# UV damage induces production of mitochondrial DNA fragments with specific length profiles

**DOI:** 10.1101/2023.11.07.566130

**Authors:** Gus Waneka, Joseph Stewart, John R. Anderson, Wentao Li, Jeffrey Wilusz, Juan Lucas Argueso, Daniel B. Sloan

## Abstract

UV light is a potent mutagen that induces bulky DNA damage in the form of cyclobutane pyrimidine dimers (CPDs). In eukaryotic cells, photodamage and other bulky lesions occurring in nuclear genomes (nucDNAs) can be repaired through nucleotide excision repair (NER), where dual incisions on both sides of a damaged site precede the removal of a single-stranded oligonucleotide containing the damage. Mitochondrial genomes (mtDNAs) are also susceptible to damage from UV light, but current views hold that the only way to eliminate bulky DNA damage in mtDNAs is through mtDNA degradation. Damage-containing oligonucleotides excised during NER can be captured with anti-damage antibodies and sequenced (XR-seq) to produce high resolution maps of active repair locations following UV exposure. We analyzed previously published datasets from *Arabidopsis thaliana, Saccharomyces cerevisiae*, and *Drosophila melanogaster* to identify reads originating from the mtDNA (and plastid genome in *A. thaliana*). In *A. thaliana* and *S. cerevisiae*, the mtDNA-mapping reads have unique length distributions compared to the nuclear-mapping reads. The dominant fragment size was 26 nt in *S. cerevisiae* and 28 nt in *A. thaliana* with distinct secondary peaks occurring in 2-nt (*S. cerevisiae*) or 4-nt (*A. thaliana*) intervals. These reads also show a nonrandom distribution of di-pyrimidines (the substrate for CPD formation) with TT enrichment at positions 7-8 of the reads. Therefore, UV damage to mtDNA appears to result in production of DNA fragments of characteristic lengths and positions relative to the damaged location. We hypothesize that these fragments may reflect the outcome of a previously uncharacterized mechanism of NER-like repair in mitochondria or a programmed mtDNA degradation pathway.

## INTRODUCTION

Mitochondria are vital organelles involved in energy production and cellular metabolism. Due to the endosymbiotic (alphaproteobacterial) origins of mitochondria, they retain their own genomes that are replicated, repaired, and inherited independently of nuclear DNA (nucDNA). Mitochondrial genome (mtDNA) mutation rates show over a 4000-fold variation across eukaryotes (1–4), which likely reflects a wide range of mtDNA replication and repair mechanisms. However, significant gaps in our understanding of mtDNA repair mechanisms still remain (5).

The existence of multiple mtDNA copies within a cell (6) led to a historical hypothesis that DNA repair mechanisms might not be necessary because damaged mtDNA could be degraded without undergoing repair and undamaged mtDNA could act as a template for mtDNA synthesis (7, 8). This idea was bolstered by the observation in metazoans that mtDNA mutation rates are much higher than nucDNA mutation rates (1) and the fact that mitochondria are an abundant source of DNA damaging reactive oxygen species (9, 10). In subsequent decades, however, researchers have determined that mtDNA repair is an important component of mtDNA maintenance and have begun to work out the mechanisms of various mtDNA repair pathways (11).

With only one known exception (12), mtDNA repair enzymes are encoded in the nucDNA, translated in the cytosol, and targeted to the mitochondria (13, 14). In some cases, mtDNA repair pathways are highly similar to nucDNA repair pathways, often utilizing enzymatic machinery that is dual-targeted to the nucleus and the mitochondria (15). For example, chemically modified mtDNA and nucDNA bases are both removed through base excision repair (BER), which is perhaps the most ubiquitous and best studied mtDNA repair pathway (16). In contrast, mtDNAs appear to lack canonical mismatch repair (MMR), the principal pathway for correcting mismatches that arise through erroneous base incorporation during DNA replication in nucDNA (17). Instead, various novel/non-canonical mismatch repair pathways may fill this role, with a piecemeal, taxon-specific distribution. For example, the Y-box binding protein YB-1 has been shown to play a role in mismatch elimination in human cell lines, primarily through mismatch recognition and binding (18). Meanwhile, plants appear to utilize a non-canonical mismatch repair pathway reliant on homologous recombination, facilitated by *MSH1*, a gene with evolutionary signatures of horizontal transfer between plants and giant viruses (19).

Nucleotide excision repair (NER) is the major nucDNA repair pathway for repair of bulky DNA damage, a broad class of lesions that occur on one strand of DNA and are characterized by the covalent attachment of large chemical moieties or compounds (20). These bulky lesions can manifest in diverse forms as a result of the binding of various chemicals, metabolites, or environmental agents to DNA, leading to structural distortions and functional impairment. NER pathways have evolved independently in bacteria and eukaryotes, with distinct variations in the protein components and regulatory mechanisms. However, both systems follow the same general mechanism in which single-stranded incisions are made both upstream and downstream of a damaged site, followed by the removal of a damage containing oligonucleotide ranging from ∼10-13 (bacterial NER) or ∼23-30 (eukaryotic nuclear NER) nt in length. A polymerase fills the resulting gap using the opposite strand as a template, and ligation completes the NER process (21). As is the case for MMR, mtDNAs are thought to lack a conventional NER pathway. Because there are no known alternative pathways for repair of bulky DNA damage in mtDNAs, it is generally assumed that bulky mtDNA damage leads to mtDNA degradation (7, 11, 15, 22), but open questions remain regarding the molecular components of mtDNA degradation, how such degradation would be coordinated, and how new mtDNA molecules could be recovered (23, 24).

Degradation of damaged mtDNAs has been documented in metazoan and yeast cells in response to a variety of DNA damaging agents including UV (25, 26), acrolein (27), gamma irradiation (28), H_2_O_2_ (29), and enzymatically induced double-stranded breaks (30). The timelines of mtDNA degradation exhibit considerable variation depending on the organism, cell type, and DNA damaging agents involved; however, it typically proceeds slowly (taking as long as 72 hours in some cases; Bess *et al.* 2012, 2013). MtDNA degradation is frequently associated with mitochondrial fission and mitochondrial-specific autophagy, known as mitophagy (27, 28). Mitophagy increases during genotoxic stress, but it also occurs in unperturbed cells as part of normal mitochondrial turnover and cellular energetics (31), and defects in mitophagy are associated with multiple human diseases (32, 33).

UV light is a potent mutagen capable of causing multiple bulky lesions, predominately in the form of cyclobutane pyrimidine dimers (CPDs; ∼80% occurrence) but also as pyrimidine– pyrimidone (6–4) photoproducts ((6–4)-PPs; ∼20% occurrence) (34). In addition to repair through NER, some organisms possess photolyases for the direct chemical reversal of photodamage. Photolyases are damage-specific, meaning a CPD photolyase can only repair CPDs and (6-4)PP photolyases can only repair (6-4)PPs. All photolyases use blue light as an energy source, and they tend to have a spotty distribution across the tree of life. Roughly half of bacteria, a quarter of archaea, most plants and fungi, and most vertebrates possess CPD photolyases; (6-4)PP photolyases are generally not as common (35–37). Photolyases have also been shown to repair photodamage in mtDNAs of some plants (38) and some fungi (22, 39). For other groups, such as mammals, there is no known mechanism for the repair of photodamage in mtDNA.

A handful of studies aimed at detecting NER in mtDNA have yielded negative results (7, 22, 38, 40–42). The earliest experiments leveraged the CPD nicking T4 endonuclease V to measure the amount of CPDs in mtDNAs of UV exposed cells. Irradiated mammalian cells given time for dark repair (NER) or light repair (photolyase) showed the same amount of mtDNA CPDs as irradiated cells given no time for repair, suggesting there is a complete lack of photodamage repair in mammalian mtDNA (7). Similar studies found that the yeast *Saccharomyces cerevisiae* also lacks dark repair of CPDs in mtDNA but does exhibit light repair (22, 40), and subsequent work established that a dual-targeted CPD photolyase protects both nuclear and mitochondrial DNA in *S. cerevisiae* (39). Tests for NER in mtDNA using qPCR in rice (38) and zebrafish (41) found no reduction in the number of polymerase blocking lesions after irradiated organisms were given periods of dark repair. qPCR studies with mice cells did detect a decrease in frequency of polymerase blocking lesions in mtDNA after long periods of repair (8 to 24 hours), but this was attributed to the repair of non-pyrimidine dimer polymerase blocking lesions, which can also be induced through UV irradiation (43). It therefore remains unclear if and how eukaryotes repair pyrimidine dimers in mtDNA. While photolyases may fill this role for some eukaryotes, they are missing entirely in some groups (mammals) or are only partially represented, such as in *S. cerevisiae*, which lacks a photolyase for the repair of the (6-4)PPs (Sancar 2004).

In recent years, a series of DNA sequencing techniques leveraging antibodies that specifically recognize CPDs or (6-4)PPs have been developed to characterize pyrimidine dimer formation and repair on genome-wide scales (45–48). One technique called DDIP-seq uses anti-damage antibodies to capture and sequence damage-containing molecules from samples of sonicated DNA (∼100 to 300 bp) (49). A DDIP-seq study with human HaCaT cells (keratinocyte cell line) and anti-CPD antibodies showed that CPD damage occurs at a high rate in mtDNA immediately following UV exposure. Surprisingly, after 24 hours allowing for repair, as much as 50% of the mtDNA damage had disappeared (47), contrasting with previous reports documenting no CPD repair in mammalian mtDNA (7, 42). Anti-damage antibodies can also be used to detect excision oligos directly in excision assays, where damage containing oligos are captured with anti-damage antibodies, 3′ radiolabeled and visualized on high-density polyacrylamide gels (50).

Another technique called XR-sequencing (XR-seq) has been particularly useful for understanding repair dynamics (48). XR-seq uses anti-damage antibodies to capture the oligonucleotides that are excised during NER (Figure 1a). These oligonucleotides are then subject to adaptor ligation, treated with photolyases, and sequenced on Illumina platforms. Sequenced reads can be aligned to reference genomes, yielding maps of active repair locations following UV exposure at single-nucleotide resolution. The technique achieves an extremely high sensitivity through the combined action of multiple filtering steps built into the library preparation (51). First, the antibodies have a high specificity for their damage targets (52), as evidenced by control immunoprecipitations with unirradiated cells, which yield no detectable DNA on polyacrylamide gels (53). The anti-CPD and anti-(6-4)PP antibodies may bind damage in both ssDNA and dsDNA (52). However, dsDNAs containing CPDs should not receive adaptors, which anneal to ssDNA through overhanging, random 5-nt sequences. In addition, the photolyases used to reverse the damage have very stringent ssDNA activity (54), as evidenced by the lack of amplification in control libraries not treated with photolyases (53). XR-seq experiments have been performed with cells or tissue samples from *Homo sapiens* (48), *Mus musculus* (55), *Microcebus murinus* (56), *Drosophila melanogaster* (57), *S. cerevisiae* (58), and *Arabidopsis thaliana* (53). The mtDNA-mapping reads from these datasets remain largely unexplored.

**Figure 1.**
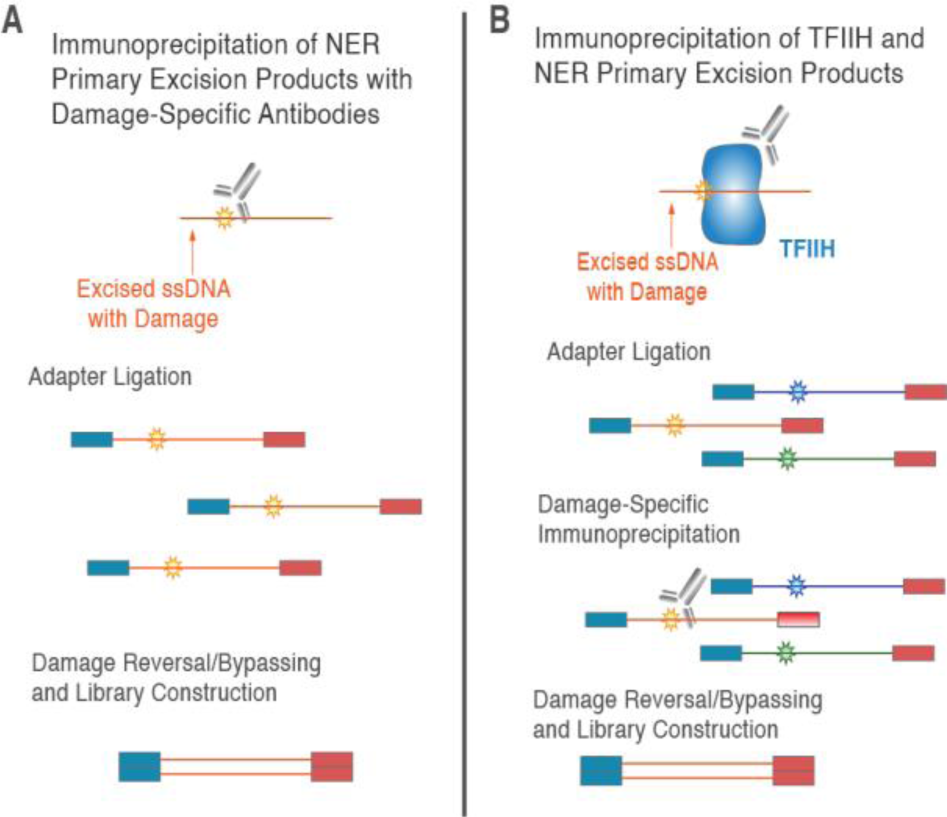
Overview of XR-seq protocol. Panel A shows direct capture of damage containing excised oligomers as performed in experiments with S. cerevisiae, A. thaliana, and D. melanogaster. After immunoprecipitation with damage specific antibodies, adaptors are attached to excised and the damaged sites are repaired by a photolyase before the molecules are amplified and sequenced. Panel B shows the alternative XR-seq approach, which includes an initial immunoprecipitation against TFIIH (the enzymatic complex that associates with excised oligomers in mammalian cells). Then, adaptors are ligated to the ssDNA fragments before a second immunoprecipitation with anti-damage antibodies, photolyase damage reversal, and library amplification/sequencing.

It is possible that previous attempts to detect NER in mtDNA may have failed because of a relatively weak signal of mtDNA repair compared to dominant signal of NER in nucDNA (59). We reasoned that the high sensitivity of XR-seq would provide increased power for detecting a NER or NER-like pathway active in mtDNA. If there is no NER or NER-like pathway for excision of photodamage or other bulky DNA lesions in mtDNA (as is generally thought) and instead such lesions lead to mtDNA degradation and turnover, the XR-seq data can still provide valuable insights into fate of photodamage during degradation and whether degradation is ordered or localized to certain regions of the genome. Published mammalian XR-seq datasets are unsuitable for such mtDNA analysis because they include an initial immunoprecipitation against TFIIH, a nuclear-localized protein complex that associates with excised oligonucleotides in mammalian NER (Figure 1b) (60). Therefore, in this study, we analyzed the mtDNA-mapping reads from published *S. cerevisiae*, *A. thaliana* and *D. melanogaster* datasets, in which the extracted small DNA molecules were immediately immunoprecipitated with anti-damage antibodies (anti-CPD or anti-(6-4)PP) without an initial TFIIH immunoprecipitation (Figure 1a).

## METHODS

### XR-seq datasets

The XR-seq datasets from *S. cerevisiae, A. thaliana* and *D. melanogaster* were generated in previous experiments (Li *et al.* 2018; Oztas *et al.* 2018; Deger *et al.* 2019; respectively). The methods used to generate those data sets are briefly summarized here. In the *A. thaliana* experiment, plants were irradiated with 120 J/m^2^ UVC at eight different times (spaced 3 hours apart) throughout a 24 hour day-night cycle and given 30 minutes of ‘dark repair’ time (53). In the *S. cerevisiae* experiment, cells were grown to late log phase and then irradiated with 120 J/m^2^ UVC and given either 5, 20 or 60 minutes of ‘dark repair’ time (58). In the *D. melanogaster* experiment, S2-DGRC cells were grown to 25-80 % confluence and then irradiated with 20 J/m^2^ UVC and given either 0.16, 0.5, 8, 16 or 24 hours of ‘dark repair’ time (57). In all three experiments, two biological replicates were included for each timepoint. The library preparation protocols were similar in all experiments, though there were differences in the methods of DNA extraction. Specifically, for *S. cerevisiae* and *D. melanogaster,* cells were disrupted through bead beating and the excised DNA was enriched by Hirt lysis, where salt is used to precipitate away the chromatin fraction of the cell lysate, and through G-50 column filtration, which further depletes the chromatin fraction (51, 57, 58). For *A. thaliana*, whole leaves were frozen in liquid nitrogen and ground into a powder before they were vortexed with glass beads (53). In all three preparations, DNA was extracted through ethanol precipitation (51). For the *S. cerevisiae* and *D. melanogaster* libraries, adaptors were added after excision products were immunoprecipitated with anti-CPD or anti-(6-4)PP antibodies (57, 58), whereas adaptor ligation preceded anti-damage immunoprecipitation for the *A. thaliana* libraries (53). In all three preparations, the adaptor-ligated products were then treated with photolyases (either CPD- or (6-4)PP-specific, depending on the library) before the samples were amplified and sequenced using 50-nt single-read Illumina chemistry.

### Alignment

Raw XR-seq reads were downloaded from NCBI BioProject folders (*A. thaliana*: PRJNA429185, *D. melanogaster*: PRJNA577587, *S. cerevisiae*: PRJNA434118) via the SRA Toolkit fastq-dump command (ver 2.8.0; Andrews 2010). Adaptor sequences (reported in original publications: Li *et al.* 2018; Oztas *et al.* 2018; Deger *et al.* 2019) were removed with cutadapt (version 1.18; Martin 2011), using the discard untrimmed reads option. Reads were aligned to reference genomes (*A. thaliana*: TAIR10, *D. melanogaster*: dm6_UCSC, *S. cerevisiae*: sacCer3) which included the organellar genomes (*A. thaliana* mtDNA: NC_037304.1, *A. thaliana* plastid DNA (ptDNA): NC_000932.1, *D. melanogaster* mtDNA: NC_024511.2, *S. cerevisiae* mtDNA: NC_001224.1) using bowtie2 (ver 2.3.5; Langmead and Salzberg 2012) with the --phred33 flag (53).

### Alignment filtering and XR-seq analysis

Nuclear insertions of mtDNA or ptDNA (termed NUMTs and NUPTs, respectively) warrant special consideration in this analysis because repair of organelle-derived nuclear DNA through conventional NER could result in the false mapping of XR-seq reads to organelle genomes. To ensure that reads mapping to the organelle genomes truly originated from the organelle genomes, we used samtools (ver 1.9; Li *et al.* 2009) to discard reads with MAPQ scores of less than 30, effectively removing all reads which map equally well to multiple locations. As a result of this filtering step NUMTs/NUPTs which are correctly assembled in the nuclear reference (and any homologous sequences present in the assemblies) are ‘unmappable’ to either copy (organellar or nuclear). The *A. thaliana* ptDNA contains a large, inverted repeat (∼26 kb). Since both copies of the repeat would be ‘unmappable’ after filtering out reads with MAPQ scores of less than 30, we removed the second copy of the repeat (positions 128214-154478) from the reference genome and divided all read counts in the first copy of the repeat by two when calculating coverage statistics. A 641-kb NUMT on chromosome 2 of the *A. thaliana* reference genome contains more than an entire copy of the mitochondrial genome (65), which introduces a potential bias as only the identical portions of the NUMT and the mitochondrial genome will be ‘unmappable’ using a MAPQ cutoff of 30. We therefore used a modified reference where the NUMT (positions 3239038-3509765 of chromosome 2) was manually removed. While interpreting the *A. thaliana* dataset, it is therefore important to remember that some mtDNA mapping reads may be nuclear-derived. After MAPQ filtering, we used custom scripts to remove reads with mismatches (all scripts used in this study are available via https://github.com/dbsloan/mtDNA_UV_damage).

We used custom scripts to calculate the read length distributions, nucleotide frequencies and di-pyrimidine frequencies of the mtDNA mapping reads and compared them to equivalent analyses from the nuclear mapping reads, which were previously reported (53, 57, 58). We analyzed the differences in read coverage (reads per kilobase per million mapped reads; RPKM) between organellar and nuclear genomes and between different genomic regions (i.e. intergenic, intronic, protein coding (CDS), rRNA genes, and tRNA genes) of the organellar genomes.

### Excision assay

To study mtDNA-derived DNA fragments with a method independent of the XR-seq data, we performed an excision assay with *S. cerevisiae* cells exposed to UV light. To isolate mtDNA-derived DNA fragments, we produced a NER-deficient line, which in theory should be unable to produce nucDNA-derived excision oligonucleotides. Specifically, we created a deletion of the *RAD14* gene, which encodes a subunit of nucleotide excision repair factor 1 (NEF1) complex that binds to damaged DNA during NER (66). Deletions were generated through homologous recombination-mediated integration of the *NatMX4* nourseothricin resistance cassette (Goldstein and McCusker 1999) in strain FY86 (*MATα, ura3-52, leu2Δ1, his3Δ200*; Winston *et al.* 1995), which is isogenic with the S288c reference genome background. We amplified *NatMX4* from pAG25 using primers JAO2397 and JAO2398 (reported in Table S1) to generate a PCR product flanked by 42-bp homologous regions (upper case in primer sequences), targeting integration to each side of the *RAD14* ORF. We screened transformants and positively confirmed the presence of the *rad14Δ::NatMX4* deletion in two independently generated clones, using PCRs with primers flanking both sides of the insertion site (primers JAO2399 and JAO2401, reported in Table S1).

Yeast growth, UV exposure, DNA extraction, immunoprecipitation with an anti-CPD antibody, radiolabeling and DNA visualization all followed previously described protocols (51), with these exceptions; 1) UV exposure was performed in a CL-1000 UV crosslinker, which was placed on a shake plate rotating at 120 rpm to ensure even UV administration, 2) we radiolabeled the 3′ ends of the putative damaging containing DNA fragments with GTP [α-^32^P] (69) instead of ^32^P-Cordycepin due to changes in product availability, and 3) we added 5% glycerol to the 11% acrylamide gel mix and electrophoresis running buffering solutions in an attempt to reduce gel shattering while drying at 80 °C (70). Following UV exposure, all work was conducted in the dark or under yellow light to avoid the activation of photolyases. We included WT and *rad14Δ* replicates that were not exposed to UV as controls, and UV-exposed strains were given 20 minutes of repair time in YPD at 30 °C. For each of the four treatments (WT vs mutant with or without UV exposure), we included two technical replicates for a total of eight samples.

## RESULTS AND DISCUSSION

### Pre-processing of existing XR-seq datasets

We analyzed the mtDNA mapping reads from *S. cerevisiae, A. thaliana* and *D. melanogaster* XR-seq datasets to gain insights into what happens to photodamaged mtDNA. In the *A. thaliana* dataset, we also investigated ptDNA-derived reads. Due to the short length of excised oligonucleotides in NER, nuclear-derived XR-seq sequences may map incorrectly to organellar genomes during alignment. To ensure such mapping artifacts are not interpreted as organellar derived DNA fragments, we filtered our alignments to retain only uniquely mapping reads with no mismatches. We assessed the impact of this filtering step by comparing XR-seq coverage of the filtered and unfiltered alignment files and found that filtering renders 5 to 13% of organellar genomes ‘unmappable’. The fraction of each genome retained for downstream analyses, broken down by genomic region, can be found in Table 1.

**Table 1:** Fraction of each organellar genome retained after filtering to remove multi-mapping reads. Retained fractions are the averages of all replicates for each dataset.

|  | *S. cerevisiae*<br>mtDNA: CPD | *S. cerevisiae*<br>mtDNA: (6-4)PP | *A. thaliana*<br>mtDNA#: CPD | *A. thaliana*<br>ptDNA*: CPD | *D. melanogaster*<br>mtDNA: CPD |
| --- | --- | --- | --- | --- | --- |
| intergenic | 0.88 | 0.88 | 0.88 | 0.95 | 0.49 |
| intron | 0.88 | 0.88 | 0.97 | 0.97 | Not applicable |
| CDS | 0.94 | 0.94 | 0.91 | 0.96 | 0.997 |
| rRNA | 0.91 | 0.91 | 0.97 | 0.91 | 0.995 |
| tRNA | 0.97 | 0.97 | 0.61 | 0.88 | 0.9996 |
| total | 0.90 | 0.90 | 0.89 | 0.95 | 0.87 |

#Before mapping we removed a large NUMT on chromosome 2 of the *A. thaliana* nuclear genome.
\*Before mapping we removed the second copy of the large, inverted repeat (~26 kb) in the *A. thaliana* ptDNA.

### XR-seq coverage of organellar vs. nuclear genomes

We next compared the depth of XR-seq coverage (computed as reads per kilobase of mapped genome; RPKM) of the organellar and nuclear genomes (Table 2). In the *S. cerevisiae* and *A. thaliana* datasets, organellar XR-seq coverage was roughly one-third to two-fold that of the nuclear genome, while in the *D. melanogaster* data coverage of the mtDNA was over 50-fold that of the nuclear genome. Note that these estimates should not be directly interpreted as measures of the relative rates of degradation or repair in nuclear vs. organellar DNA because they do not adjust for differences in organellar genome copy per nuclear genome, a parameter known to be highly variable under different life stages (71), tissue and cell types (Herbers et al. 2019; O’Hara et al. 2019), and physiological conditions (74). The relative rates of pyrimidine dimer formation in organellar vs. nuclear DNA will also impact rates of repair, and estimates of the relative damage rates vary among species (38, 42, 43) and depend on methods of detection (75).

**Table 2:** Organellar vs. nuclear XR-seq coverage (as RPKM)

|  | *S. cerevisiae*<br>mtDNA: CPD | *S. cerevisiae*<br>mtDNA: (6-4)PP | *A. thaliana*<br>mtDNA: CPD | *A. thaliana*<br>ptDNA: CPD | *D. melanogaster*<br>mtDNA: CPD |
| --- | --- | --- | --- | --- | --- |
| organellar RPKM | 25.0 | 72.1 | 11.7 | 13.6 | 424.9 |
| nuclear RPKM | 82.7 | 82.3 | 8.4 | 8.4 | 6.9 |
| ratio: org/nuc RPKM | 0.30 | 0.88 | 1.39 | 1.62 | 61.58 |

### Unique length distributions of organellar-mapping reads

We next analyzed the length of XR-seq fragments mapping to organellar and nuclear genomes. For all datasets, the organellar-mapping reads contain unique length distributions compared to those mapping to the nuclear genomes. As reported in the initial publication (58), there are two peaks in the *S. cerevisiae* nucDNA mapping reads (in both anti-CPD and anti-(6-4)PP datasets), one derived from the primary excision products (23 nt) and the other (∼16 nts) presumably derived from the 5′ degradation of the primary excision products (Figures 2, S1, S6). The *S. cerevisiae* CPD and 6-4(PP) mtDNA-mapping reads show distinct peaks at read lengths of 26, 24, 22 and 20 nt (Figures 2, S2, S7). The largest mtDNA peak of 26 nt is longer than the peak length in the nuclear mapping reads of 23 nt. The *A. thaliana* mtDNA read length distributions also differ from the nucDNA read length distributions (Figures 3, S11). In the *A. thaliana* mtDNA read length distribution there is a cluster of reads 36-39 nt in length, with additional distinct peaks in read lengths of 32, 28, 24, 20 and 16 nt (Figures 3, S12). Therefore, the patterns in these datasets were similar, but the peaks were spaced at different intervals (2-nt in *S. cerevisiae* and 4-nt in *A. thaliana*).

**Figure 2.**
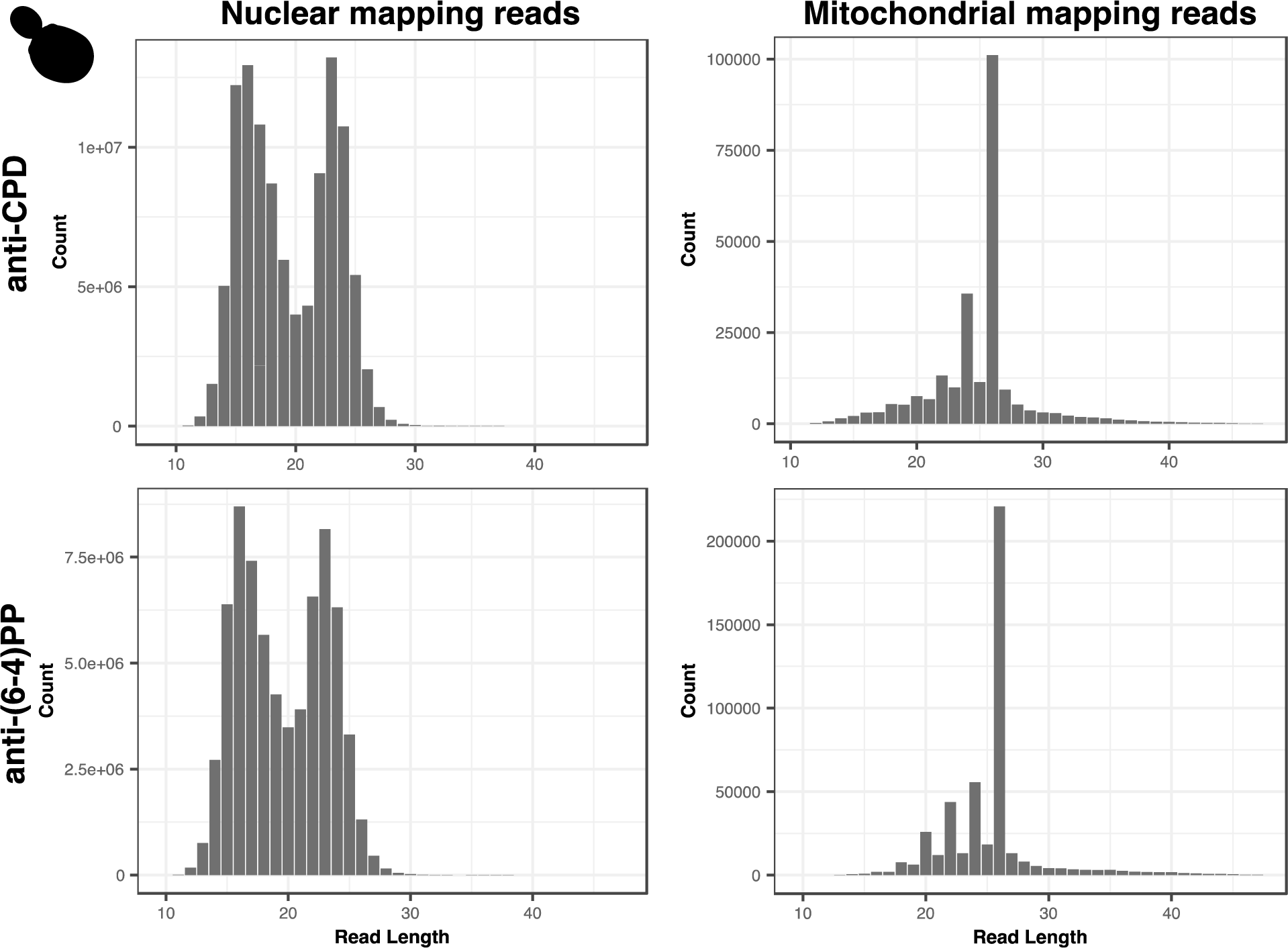
Read length distributions of nuclear and mitochondrial reads from anti-CPD and anti-(6-4)PP libraries from *S. cerevisiae.* These distributions exhibited a high degree of repeatability across samples and conditions. Pearson’s correlation analyses reveal significant correlations between the anti-CPD and anti-(6-4)PP read length distributions (R=0.9555, p=1.6E-11) as well as between anti-CPD (5 min vs 20 min; R=0.9951, p=2.2E-16) and anti-(6-4)PP (5 min vs 20 min; R=0.9765, p=3.92E-14) timepoints.

**Figure 3.**
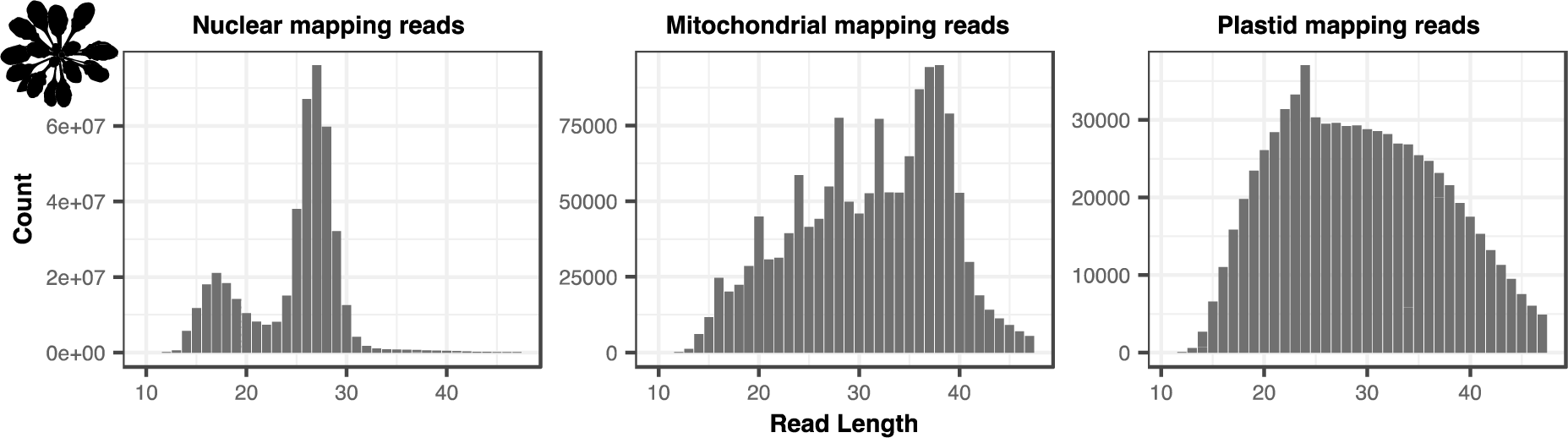
Read length distributions of nuclear, mitochondrial and plastid mapping reads from the *A. thaliana* anti-CPD libraries. Pearson’s correlation analyses reveal significant correlations between in the mtDNA read length distributions between time points (2 hours vs. 5 hours: R= 0.9899, p=2.2E-16).

The *A. thaliana* ptDNA read length distribution lacks distinct peaks occurring at regular intervals, and instead contains a single, less extreme peak comprised of reads 24 nt in length (Figures 3, S13). The *D. melanogaster* mtDNA-mapping reads have a different read length distribution compared to the nuclear-mapping reads (Figures 4, S19, S20), but the mtDNA-mapping reads lack the discrete peaks we observed in *S. cerevisiae* and *A. thaliana* organellar reads.

**Figure 4.**
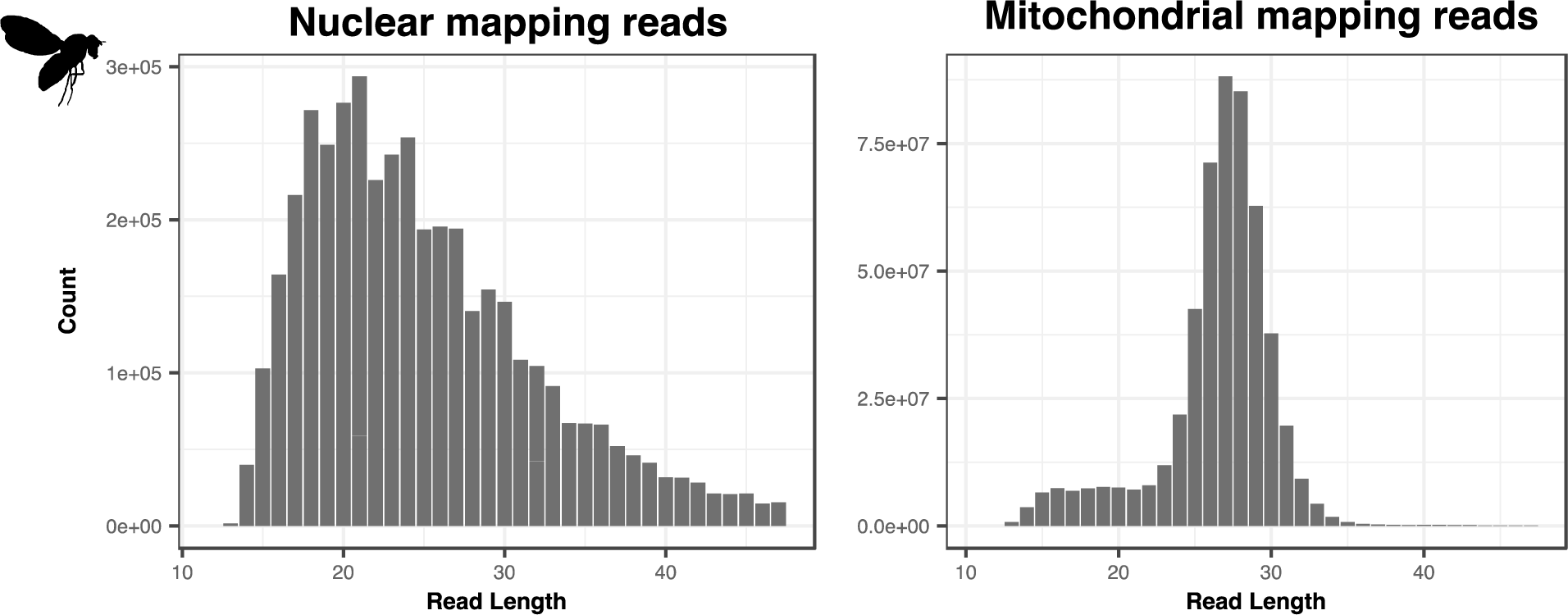
Read length distributions of nuclear and mitochondrial mapping reads from the *D. melanogaster* anti-CPD libraries.

The origins of the distinct peaks in the XR-seq read length distributions of the *S. cerevisiae* and *A. thaliana* datasets are unclear. It is possible that these DNA fragments are derived from a mitochondrial-specific NER-like pathway for photodamage repair. Alternatively, the abundance of reads of certain lengths could arise from mtDNA degradation, which would represent a previously uncharacterized mechanism of damage-induced mtDNA degradation that results in fragments of specific lengths. XR-seq experiments in *E. coli* reveal somewhat similar patterns in the sense that there are a few read lengths that account for most of the reads in the length distribution, except that in *E. coli* most reads are of 10 or 13 nt in length (76). Interestingly, there is only a single peak of 13 nt in excision assays with *E. coli* mutants lacking *UvrD*, presumably because the primary 13mer oligonucleotide is unable to dissociate from the UvrB-UvrC heterodimer without the activity of the UvrD helicase and is therefore inaccessible to the exonucleases that degrade the oligonucleotide from the 3′ end (77). In the *S. cerevisiae* nucDNA derived reads, TT peaks consistently occur 6 nt from the 3′ ends of reads, including in reads less than 23 nts (the length of the primary excision product in nucDNA NER), suggesting that nucDNA-derived reads are degraded from the 5′ ends. Given that secondary excision products have been shown to arise through exonuclease degradation of a primary oligonucleotide in *E. coli* and in the *S. cerevisiae* nucDNA NER, we hypothesize that the 26-nt peak in the *S. cerevisiae* mtDNA may be a ‘primary’ product, with the less abundant 24, 22 and 20 nt oligonucleotides arising through degradation.

Attempts to visualize and validate the read length distributions observed in XR-seq data with a conventional excision assay found that the *S. cerevisiae* mtDNA signal was undetectable above background, even when the nucDNA signal was reduced by using a nuclear NER-deficient mutant strain background (*rad14Δ*) (Figure S24). It is likely that the signal of repair or degradation from the mtDNA is relatively weak compared to the noise of the assay (see faint gray smear in every lane of Figure S24). Future efforts to identify mtDNA fragments in excision assays may benefit from increased sample volumes and from physically isolating mitochondria from the cell suspensions before immunoprecipitation with anti-damage antibodies.

### Preferential positioning of pyrimidines within organellar-mapping XR-seq reads

We analyzed the nucleotide and di-pyrimidine frequencies of all the organellar mapping reads, focusing especially on the dominant read lengths in the *S. cerevisiae* and *A. thaliana* mtDNA datasets (shown in Figure 2 and Figure 3, respectively). In the 26-nt *S. cerevisiae* mtDNA mapping reads (the most frequent length class) in the CPD dataset, adjacent thymines (TTs) are most abundant at position 7-8, with additional smaller peaks spaced at 2-nt intervals, starting at positions 10 (left panel of Figure 5; di-pyrimidine frequencies of all the (6-4)PP mapping reads including rare size classes are shown in Figure S3). The 24-nt reads show a similar TT peak pattern, though it is shifted forward two positions compared to the pattern in the 26 nt reads (i.e., a peak at position 5-6, followed by secondary peaks starting at positions 8, 10, and 12). In the 22 nt reads the TT peaks are shifted forward four positions. Therefore, the TT peaks fall in the same position when the 26, 24 and 22 nt reads are 3′ or right aligned as they are in Figure 5. In the (6-4)PP dataset, the TT peaks also fall in similar positions when the most common reads (26, 24, 22 and 20) are 3’ aligned (right panel of Figure 5, di-pyrimidine frequencies of all the (6-4)PP mapping reads including rare size classes are shown in Figure S8), though the TT peak at position 7-8 does not rise above the null expectation derived from the frequency of TTs in the *S. cerevisiae* mtDNA.

**Figure 5.**
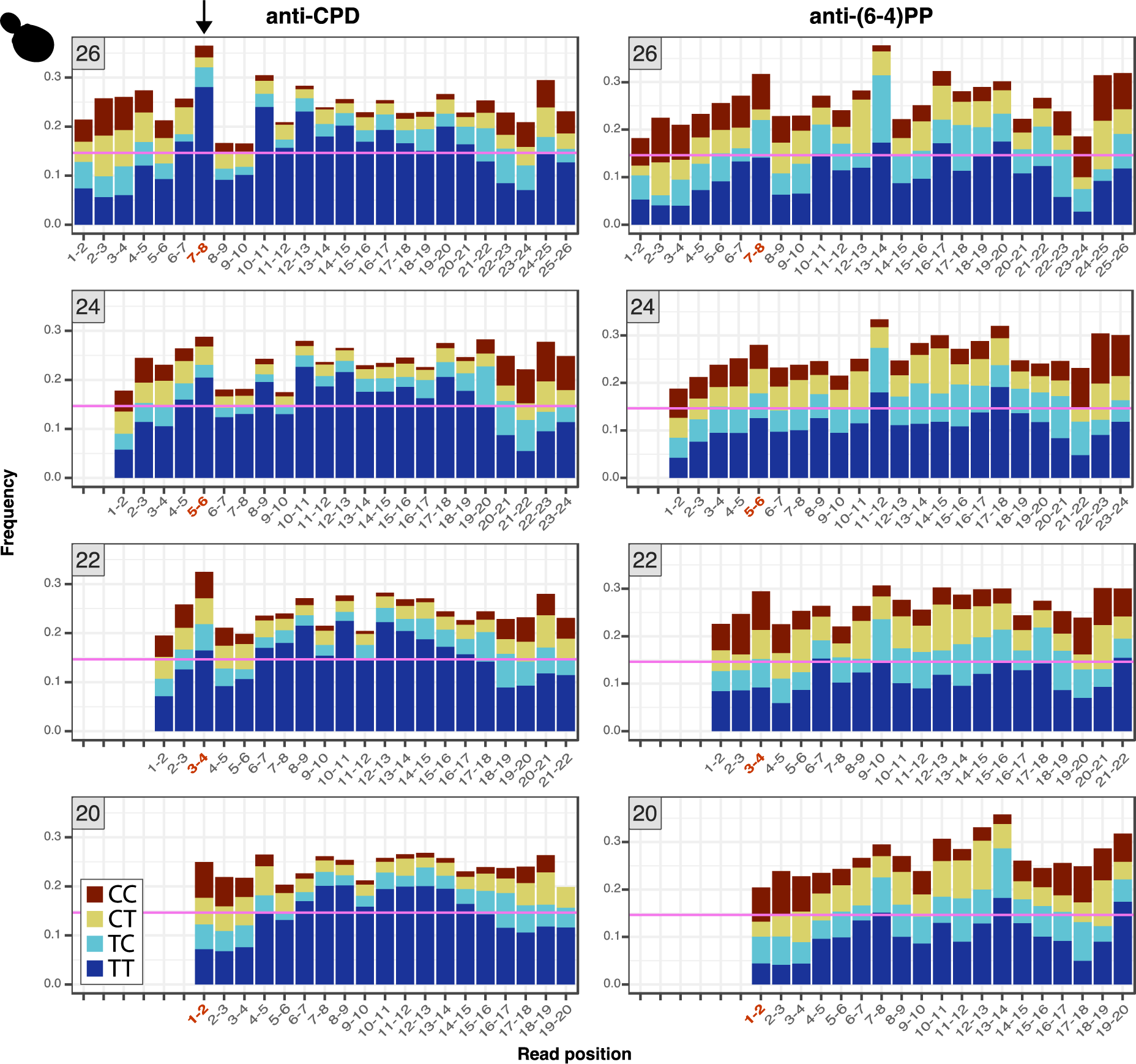
Di-pyrimidine frequencies in the most abundant read-length classes (26, 24, 22 and 20 nt) from the *S. cerevisiae* anti-CPD libraries. Read lengths are denoted in the gray boxes at the top left of each panel. The pink horizontal lines show the frequency of TT dinucleotides in the *S. cerevisiae* mtDNA, providing a null expectation for TT dinucleotide frequencies in the XR-seq reads. Positions with TT peaks in the 26 nt reads are in red, and the equivalent positions in the 3′ aligned (right aligned) 24, 22 and 20 nt reads are also in red. We approximated the 95% confidence interval as 2 times the standard error of the expected TT frequency given the number of reads included for each di-pyrimidine calculation. Given the large number of reads analyzed, 95 % confidence intervals are very small, ranging from 0.1472 ± 0.0022 for the CPD 26-nt reads to 0.1472 ± 0.0081 for the CPD 20-nt reads. As a result, all blue bars that appear above the pink line in the figure represent a significant statistical enrichment relative to the expectation and its 95% confidence interval.

The observed patterns in the mtDNA-derived XR-seq reads may arise through mtDNA degradation or through an incision-based repair process (e.g., NER-like repair). In either scenario, we propose a potential mechanism in which ‘primary’ 26-nt DNA fragments may be degraded in 2-nt intervals from the 5′ end to produce 24, 22 and 20 nt products (Figure 7, right panel). Alternatively, incisions of 6, 4 or 2 nt upstream of a CPD could yield the 26, 24 and 22 nt products, respectively (Figure 7, left panel). Under the model that these DNA fragments arise through wholesale mtDNA degradation, rather than a specific incision-based pathway, we hypothesize that TT dimers inhibit mtDNA degrading nucleases from accessing upstream or downstream nucleotides, resulting in the enrichment of di-pyrimidines at internal locations in the DNA fragments, though we are not aware of examples of exonucleases stalling at such distances from pyrimidine-dimers in the literature. Instead, previous efforts to understand the fate of excised oligonucleotides generated during NER in nucDNA have identified multiple exonucleases that can remove nucleotides up to a dimer (50, 77–79).

DNA fragments with dimers on the end are difficult to study because they can be recalcitrant to elongation by terminal transferase enzymes necessary for radiolabeling and likely to ligation of adaptors necessary for XR-seq (78). Therefore, it is possible that ‘dimer-capped’ mtDNA molecules generated through exonuclease activity up to the dimer would be undetectable in XR-seq datasets. Previous XR-seq studies have also suggested that adaptor ligation biases may drive variation in nucleotide composition within reads (80), especially at read-ends where adaptors are ligated. We cannot completely rule out the possibility that adaptor ligation biases are responsible for the enrichment of TT dinucleotides at specific positions (e.g., position 7-8 in 26-nt *S. cerevisiae* CPD reads). However, we view this as an unlikely explanation for multiple reasons: 1) the TT enrichment is internal to the fragments and not directly at the ligated ends and often outside the random 5-nt sequence used for adaptor annealing, 2) the enrichment patterns differ greatly across species and across genomes, and 3) the enriched positions shift relative to the 5′ end depending on the length of read (Figure 5).

As in the *S. cerevisiae* dataset, the TT peaks in the *A. thaliana* mtDNA mapping reads fell in the same position when the most frequent read lengths are aligned (in this case reads 32, 28, 24, 20 and 16 nt long). For the *A. thaliana* dataset, this pattern holds regardless of whether the reads are left (5′) aligned (as in Figure 6; di-pyrimidine frequencies of all the mtDNA-mapping reads including rare size classes are shown in Fig S14) or right (3′) aligned (not shown). If these DNA fragments are arising through targeted incisions, we posit that these incisions occur primarily either 6, 10, 14, 18 or 22 nt upstream of a CPD, and either 8, 12, 16, 20 or 24 nt downstream of a CPD. Alternatively, the 4-nt spacing of pyrimidine dimers could be explained by regular degradation of a primary excision product of undetermined length. Yet another possibility might be that if these fragments are arising through mtDNA degradation, it would again appear that degrading nucleases are unable to access DNA within a certain distance of pyrimidine dimers.

**Figure 6.**
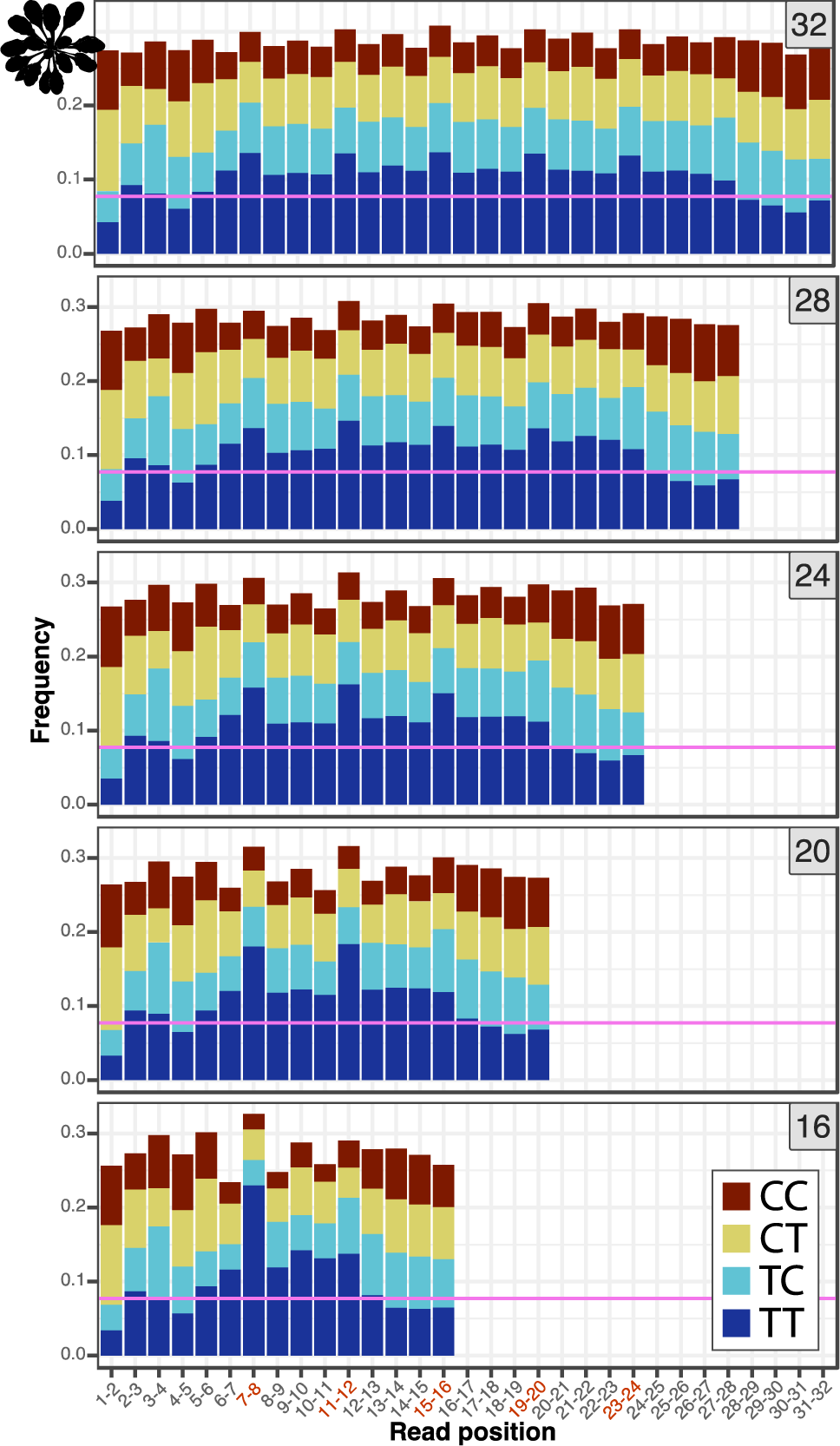
Di-pyrimidine frequencies in the most abundant read-length classes (32, 28, 24, 20 and 16 nt) from the *A. thaliana* mtDNA mapping reads. Read lengths are denoted in the gray boxes at the top right of each panel. The pink horizontal lines show the frequency of TT dinucleotides in the *A. thaliana* mtDNA, providing a null expectation for TT dinucleotide frequencies in the XR-seq reads. See Figure 5 for a description of calculating 95% confidence intervals around this expectation. These confidence intervals were very small due to the large number of reads, ranging from 0.0743 ± 0.0033 for the 16 nt reads to 0.0743 ± 0.0018 for the 32 nt reads.

**Figure 7.**
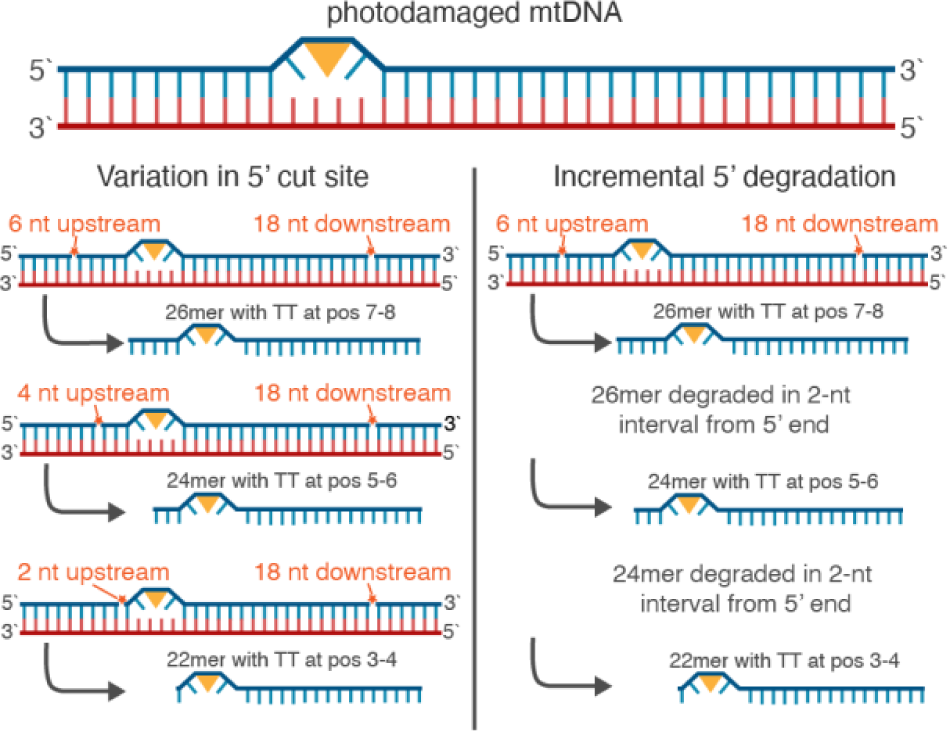
Proposed alternative explanations for unique read length distributions and dinucleotide composition patterns in the *S. cerevisiae* anti-CPD and anti (6-4)PP datasets.

The *A. thaliana* ptDNA and *D. melanogaster* mtDNA reads lack obvious di-pyrimidine patterns (Figures S15, S21). Interestingly, the *D. melanogaster* mtDNA mapping reads show extreme nucleotide biases at both the 5′ and 3′ ends of reads (Figure S22). Such biases may be driven by biased composition of the overhanging Ns that allow for adaptor annealing. However, nuclear mapping XR-seq reads do not display extreme nucleotide biases at read ends (see Figure 1c; Deger et al. 2019). End biases are also mostly absent from the *S. cerevisiae* and *A. thaliana* organellar mapping reads (Figures S4, S9, S16, S17; respectively), which were created with the same or similar adaptors. Another explanation could be that in the *D. melanogaster* mtDNA, distance from a CPD is not important in determining upstream or downstream incision sites, and instead local sequence contexts drive incision locations. Such a phenomenon would also explain why the *D. melanogaster* read length distribution lacks discrete peaks (Figure S20) and why the *D. melanogaster* mtDNA-mapping reads lack an enriched localization of di-pyrimidines (Figure S21).

### Variation in the distribution of XR-seq reads among genomic regions

We determined the location of the organellar mapping reads as either intergenic, CDS (protein coding), intronic, tRNA coding, or rRNA coding. In both *S. cerevisiae* datasets (CPD and (6-4)PP), we find elevated coverage of genic regions (CDS, rRNA and tRNA) compared to coverage in intergenic regions (Figures S5, S10; left panel). This pattern is consistent with trends in the *S. cerevisiae* nuclear genome (58), where increased genic XR-seq coverage is attributed to transcription-coupled NER (TC-NER). Another feature of TC-NER is increased coverage of the template DNA strand compared to the coding DNA strand. In both *S. cerevisiae* datasets (CPD and (6-4)PP), we find elevated coverage of the template strand compared to the coding strand in intronic, tRNA and rRNA regions of the genome, consistent with TC-NER-like processes in these regions (Figures S5, S10; right panel). However, CDS regions show slightly elevated coverage of the coding strand compared to the template strand, which is inconsistent with expectations of TC-NER. Importantly, differences in XR-seq coverage between genomic regions may also arise from different levels of pyrimidine dimer formation, which has been shown to vary across nucDNAs due to variation in local sequence motifs and nucleosome density (45). Organellar DNA lacks nucleosomes and is instead packaged in nucleoids, which can vary in protein components based on developmental and physiological status of a given organelle, but are generally assumed to confer many of the same protective benefits as nucleosomes (23, 24, 81).

In the *A. thaliana* mtDNA, we see slightly elevated XR-seq coverage of the CDS compared to the intergenic regions of the genome, but rRNA and tRNA genes, which are typically expressed more highly than CDS regions (82, 83), have XR-seq coverage below or near the level of intergenic sequence (Figure S18, top left panel). This suggests that increases in expression may not correlate with increased levels of of incisions or repair activity as is observed in the *A. thaliana* nucDNA due to TC-NER (53). In the *A. thaliana* ptDNA, we see relatively even levels of CDS and intergenic coverage, but decreased coverage of rRNAs and tRNAs (Figure S18, top right panel), again opposite of the expectations under a TC-NER-like repair model where more highly expressed genes receive increased NER protection. If the organellar-derived DNA fragments arise through organellar genome degradation rather than by NER-like pathways, variation in XR-seq read depth across genomic compartments may provide a snapshot of variation in damage formation. There are no large effect asymmetries in coding vs template strand in the *A. thaliana* data (Figure S18, bottom panels) except for in ptDNA rRNA genes, where template coverage is roughly 2-fold that of the coding strand. It is difficult to know whether these asymmetries arise through variation in damage formation, NER-like repair, or asymmetrical DNA degradation.

In the *D. melanogaster* mtDNA, we find a drastic reduction in coverage of the intergenic portion of the genome compared to the CDS, rRNA and tRNA genes (Figure S23). Metazoan mtDNAs are extremely gene dense, so essentially all of the ‘intergenic’ sequence in the *D. melanogaster* mtDNA is located in the AT-rich region of the genome, which serves as the mtDNA replication origin and termination sites. Given the preponderance of thymines in this region, one might expect an increase in CPD formation compared to other regions of the genome, making the lack of XR-seq read in this region intriguing. However, AT rich sequences also experience negative amplification biases during the PCR stages of library construction (84–86), so comparisons of XR-seq coverage between regions of varied AT content must be made cautiously.

## CONCLUSION

Early studies that found no repair of UV-damage mtDNAs in human and yeast cells (7, 22) helped shape the notion that mitochondria lack DNA repair altogether and that damaged mtDNA molecules are simply degraded, with undamaged copies serving as templates for new mtDNA synthesis (8). While subsequent investigations have unveiled that specific types of mtDNA base damage such as deamination, simple alkylation, and oxidation can indeed be effectively repaired within the mitochondria, it is still generally accepted that all eukaryotes lack any NER-like pathway for repair of bulky DNA damage in mtDNAs (15, 87–89). MtDNA damage has been demonstrated to lead to mtDNA degradation in a variety of instances (25–30), but this process remains enigmatic, with open questions as to how damaged mtDNAs are distinguished from healthy mtDNAs, how damaged mtDNAs promote fusion and or mitophagy (25, 27, 33), and which enzymes actually degrade the mtDNA (30, 90, 91).

Our analysis of XR-seq experiments from diverse eukaryotes shows that mitochondrially derived DNA fragments of characteristic length and nucleotide composition are produced following mtDNA photodamage. As we have laid out, we envision two potential mechanisms that could be responsible for productions of these DNA fragments: 1) an NER-like repair pathway functioning in mitochondria, or 2) a previously uncharacterized programmed degradation of damaged mtDNA. Either of these possibilities point to the exciting prospect of novel maintenance or processing in response to exogenous damage. A key next step in differentiating between these and other possible models will be to be identify the specific molecular machinery that produces the observed DNA fragments in response to UV damage.

## ACKNOWLEDGEMENTS

We thank Luis Brieba, José Gualberto, and Ogün Adebali for helpful discussions about these results. Gus Waneka and Dan Sloan were supported by the National Institutes of Health (NIGMS R35GM148134). Wentao Li was supported by National Institute of Environmental Health Sciences (NIEHS, R00 ES030015). Genome stability research in the Argueso laboratory is supported by National Institutes of Health award R35GM11978801

---

**Figure S1.**
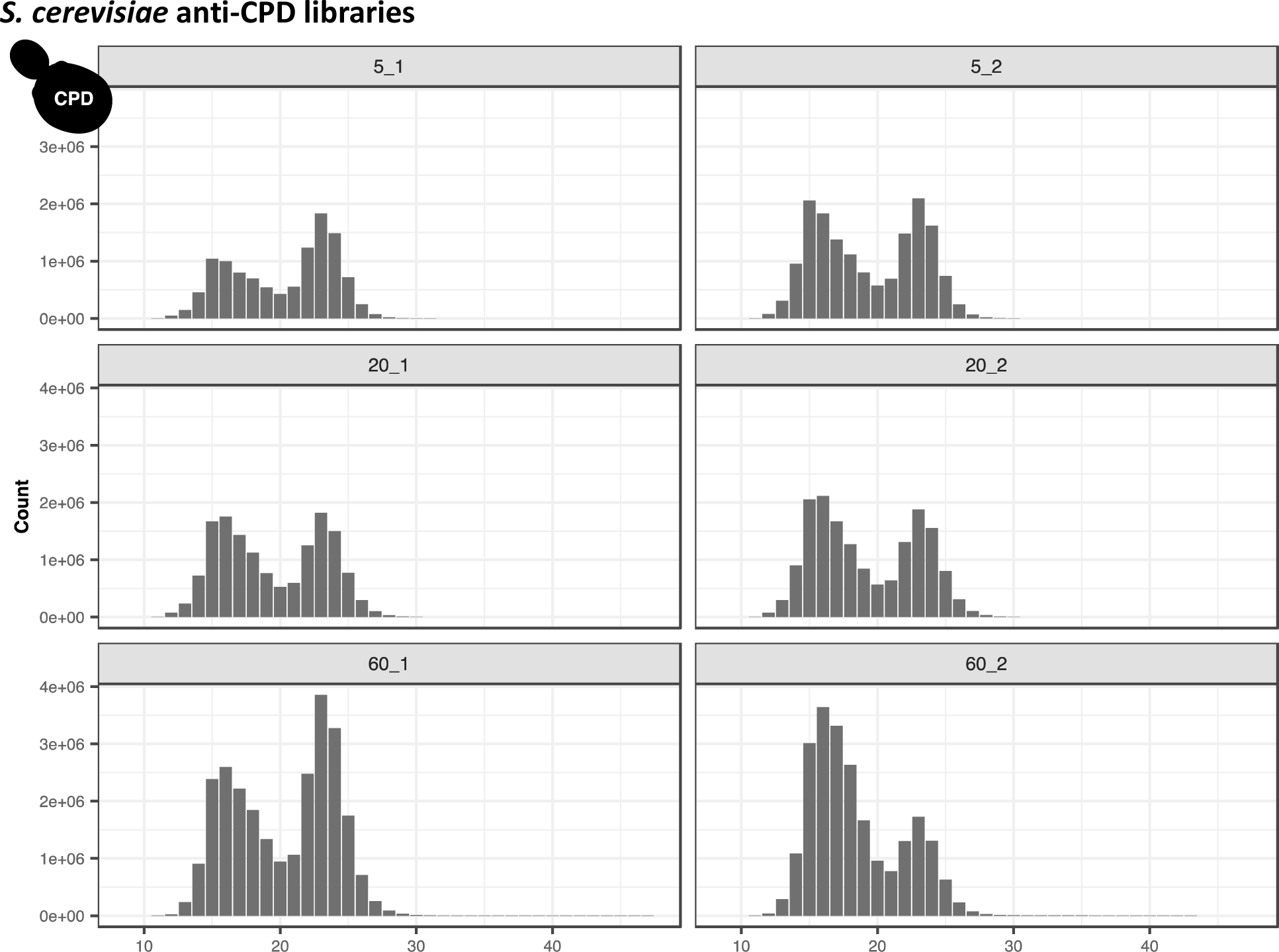
Length distributions of the nuclear-mapping reads from the *S. cerevisiae* anti-CPD libraries. The panel labels show the library ID, with the first number indicating the repair time (5, 20 or 60 minutes) and the second number indicating the replicate number (1 or 2).

**Figure S2.**
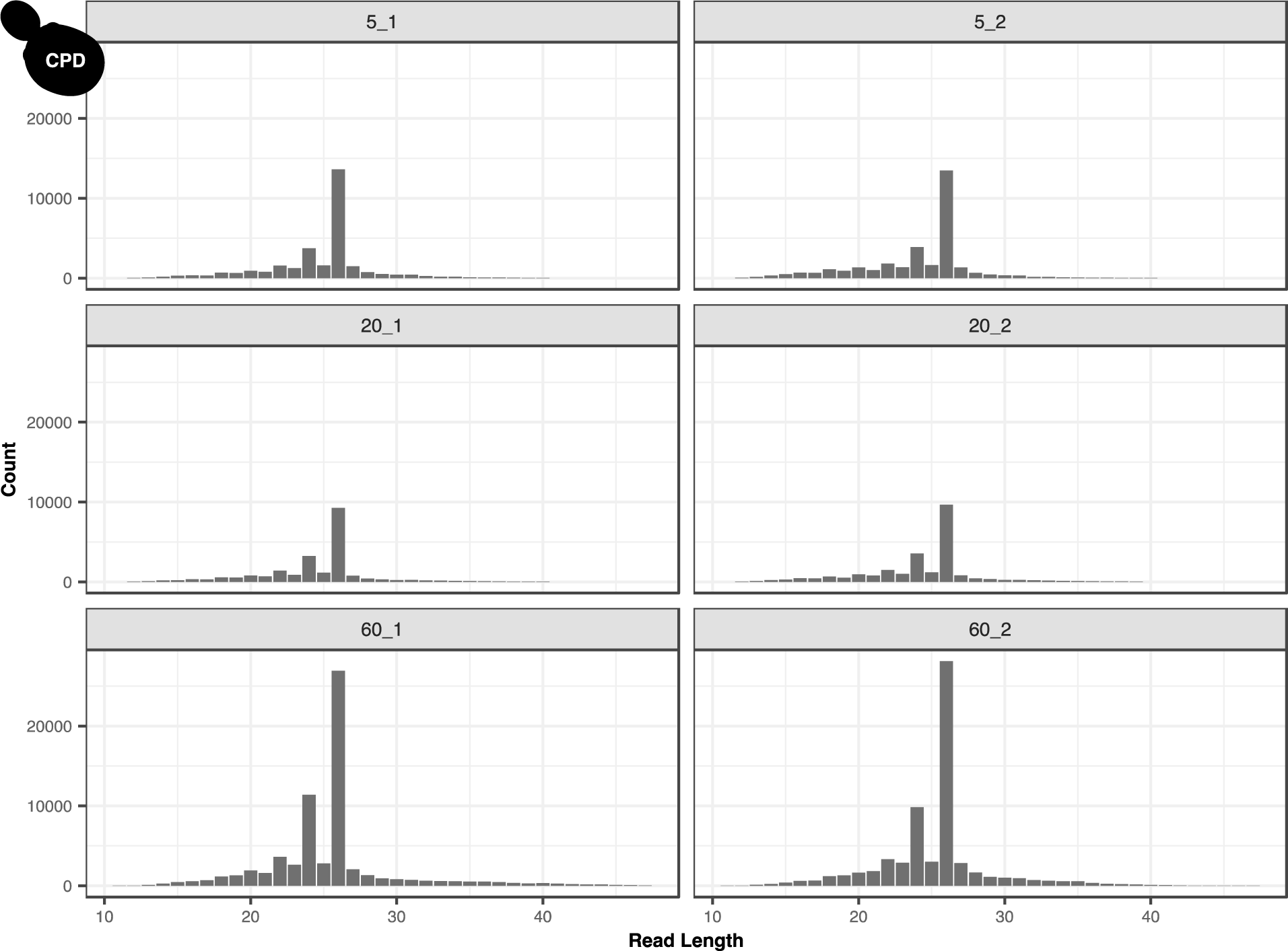
Length distributions of the mtDNA mapping reads from the *S. cerevisiae* anti-CPD libraries. The panel labels show the library ID, with the first number indicating the repair time (5, 20 or 60 minutes) and the second number indicating the replicate number (1 or 2).

**Figure S3.**
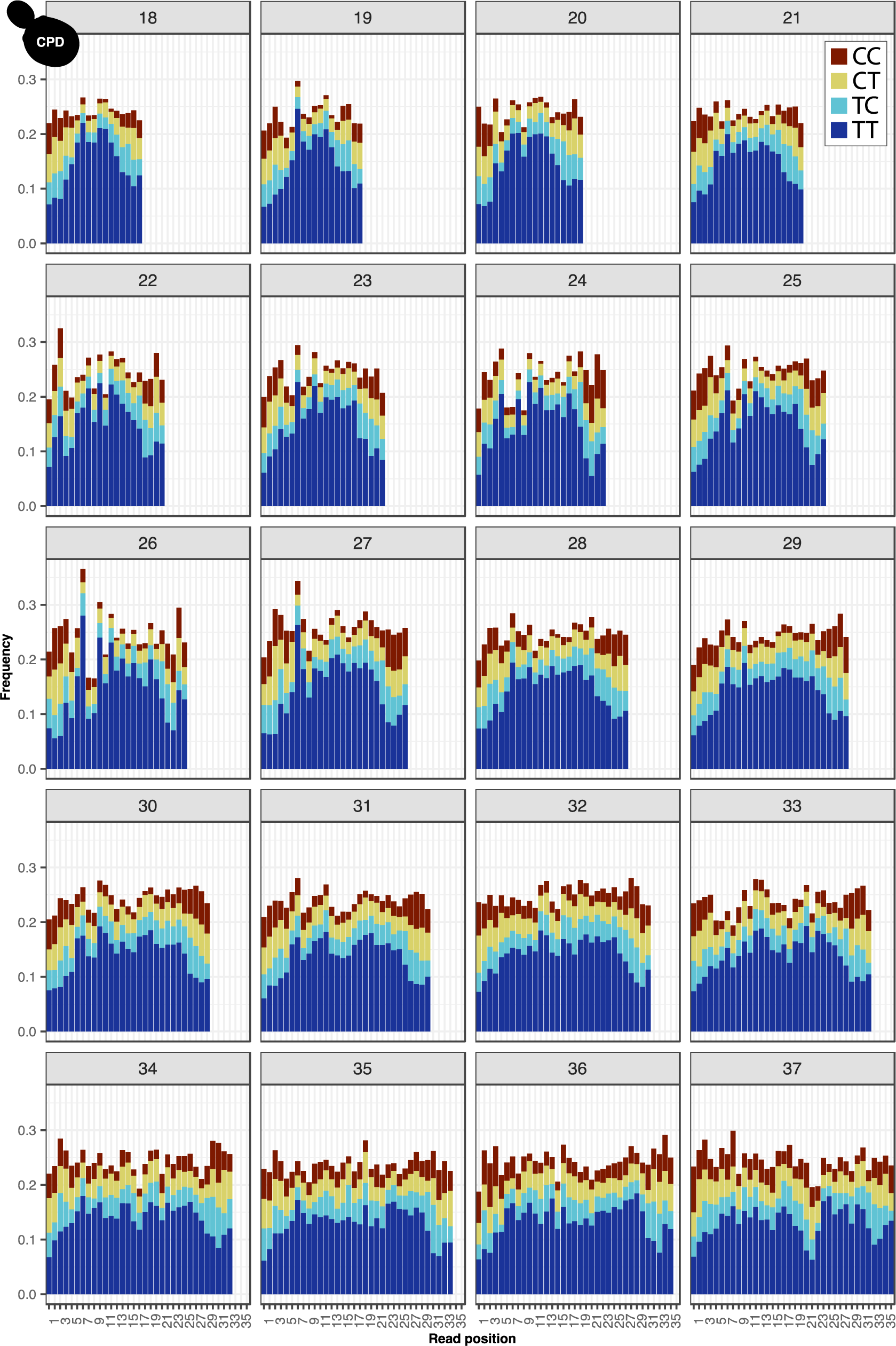
Di-pyrimidine frequencies in the mtDNA mapping reads from the *S. cerevisiae* anti-CPD libraries for all read-length classes.

**Figure S4.**
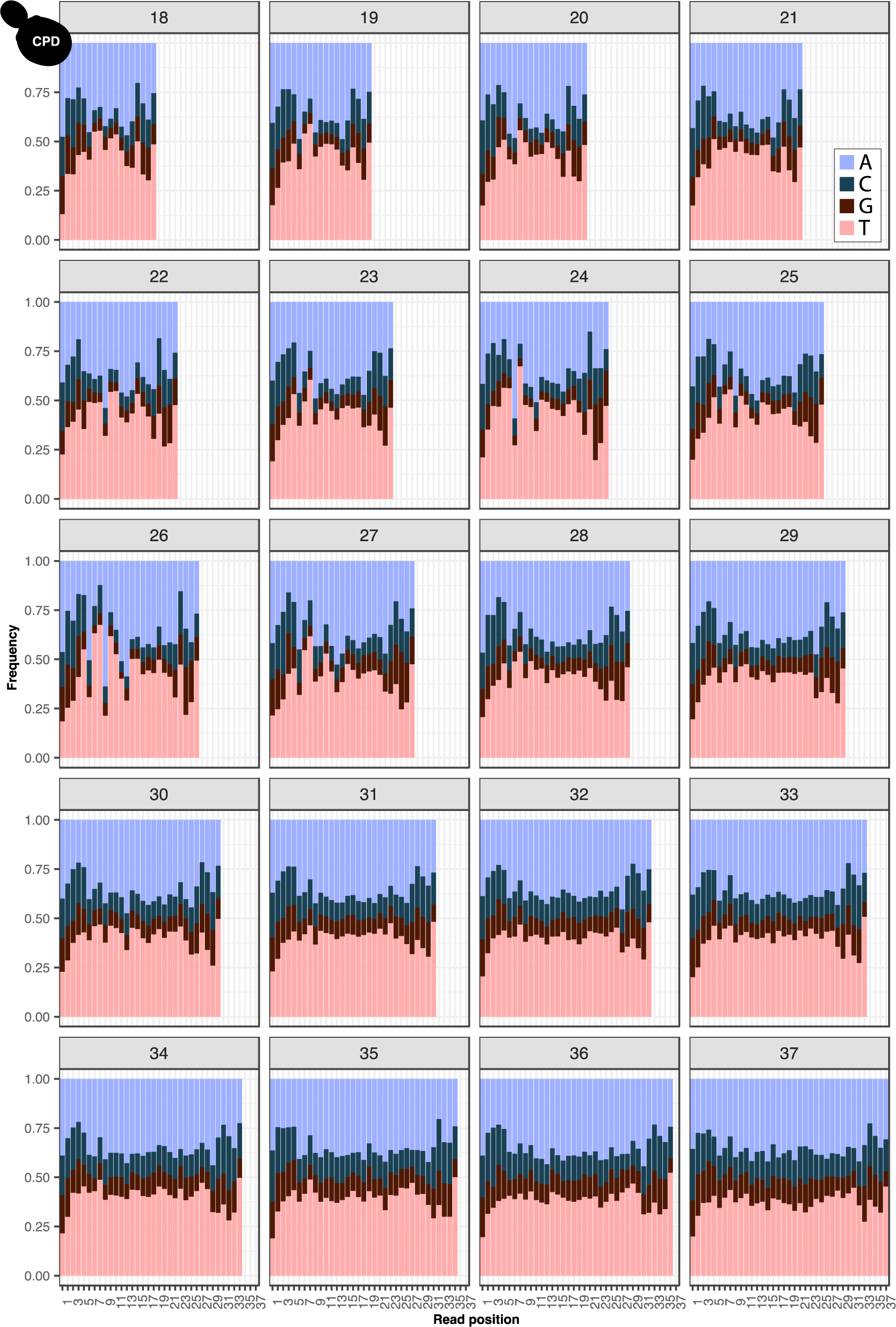
Nucleotide frequencies in the mtDNA mapping reads from the *S. cerevisiae* anti-CPD libraries for all read-length classes.

**Figure S5.**
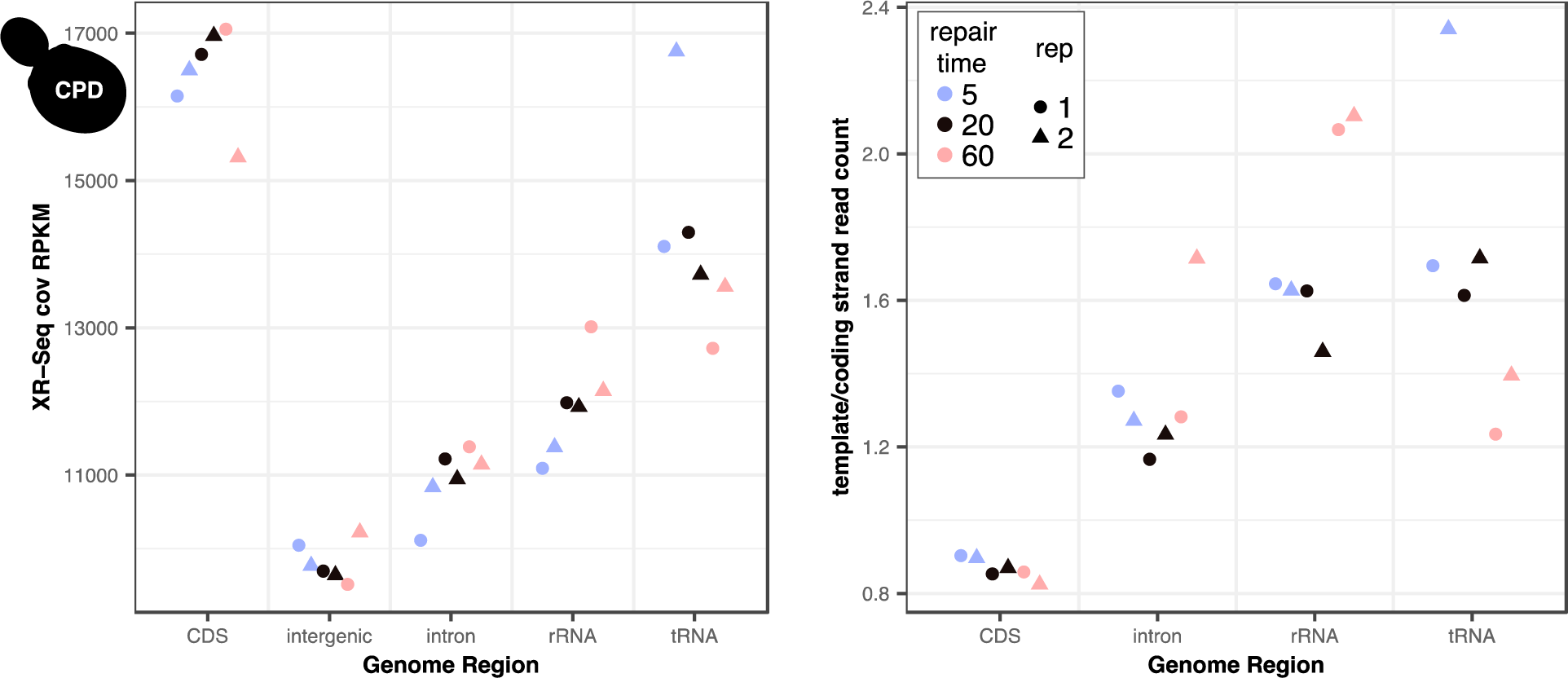
Depth of coverage (RPKM) and strand assymetry by genomic region in the mtDNA-mapping reads from the *S. cerevisiae* anti-CPD libraries.

**Figure S6.**
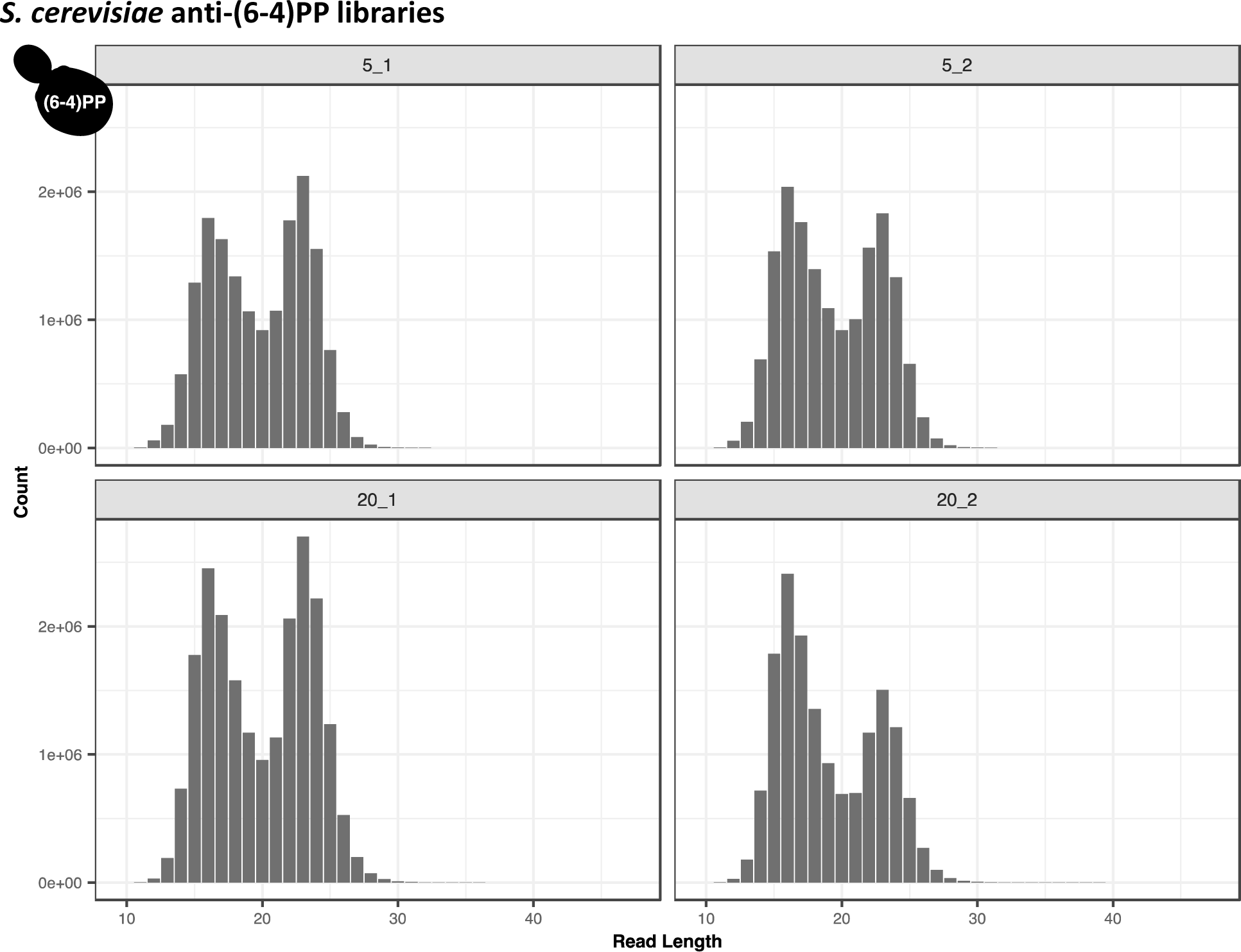
Length distributions of the nuclear-mapping reads from the *S. cerevisiae* anti-(6-4)PP libraries. The panel labels show the library ID, with the first number indicating the repair time (5 or 20 minutes) and the second number indicating the replicate number (1 or 2).

**Figure S7.**
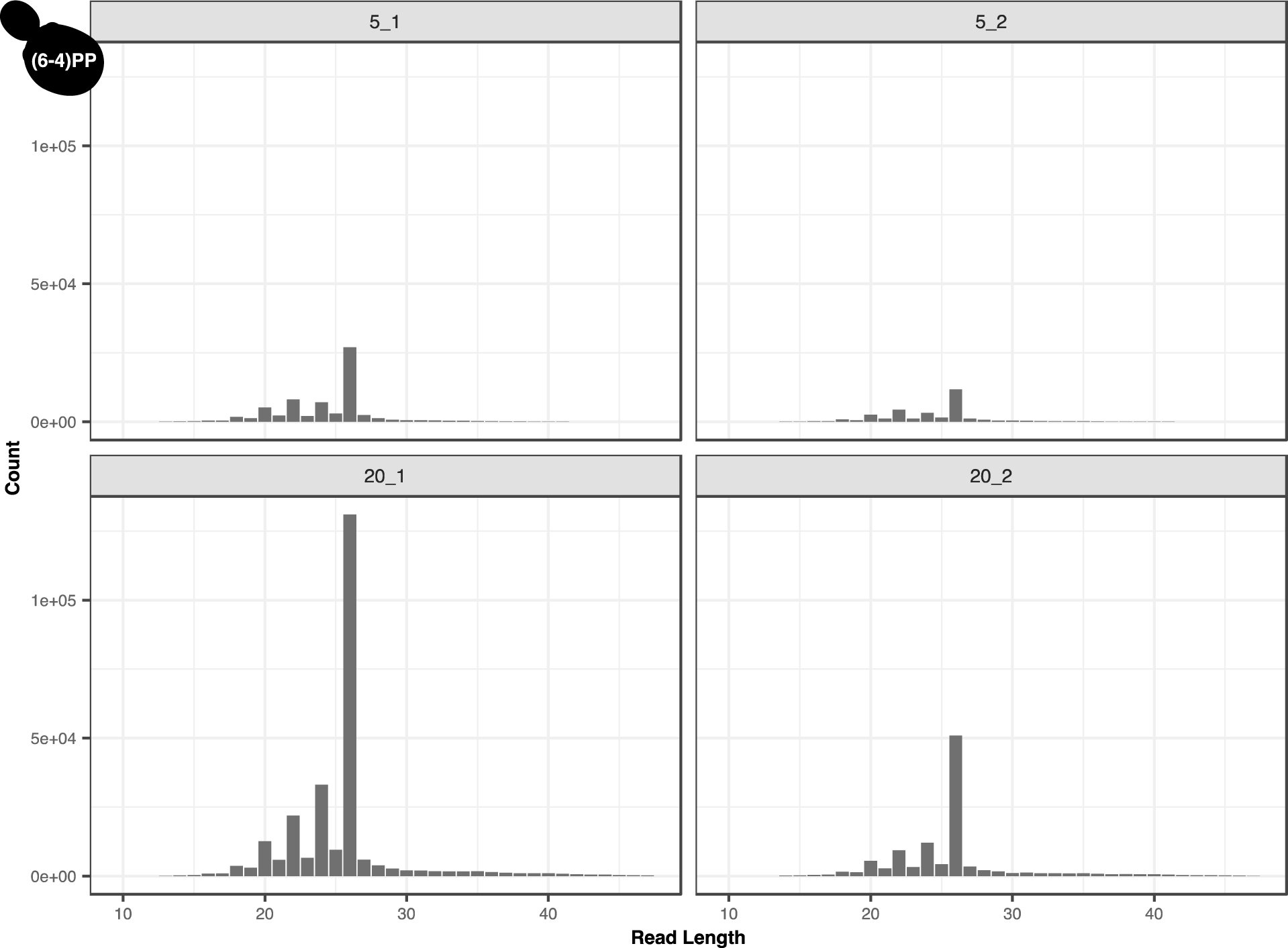
Length distributions of the mtDNA-mapping reads from the *S. cerevisiae* anti-(6-4)PP libraries. The panel labels show the library ID, with the first number indicating the repair time (5 or 20 minutes) and the second number indicating the replicate number (1 or 2).

**Figure S8.**
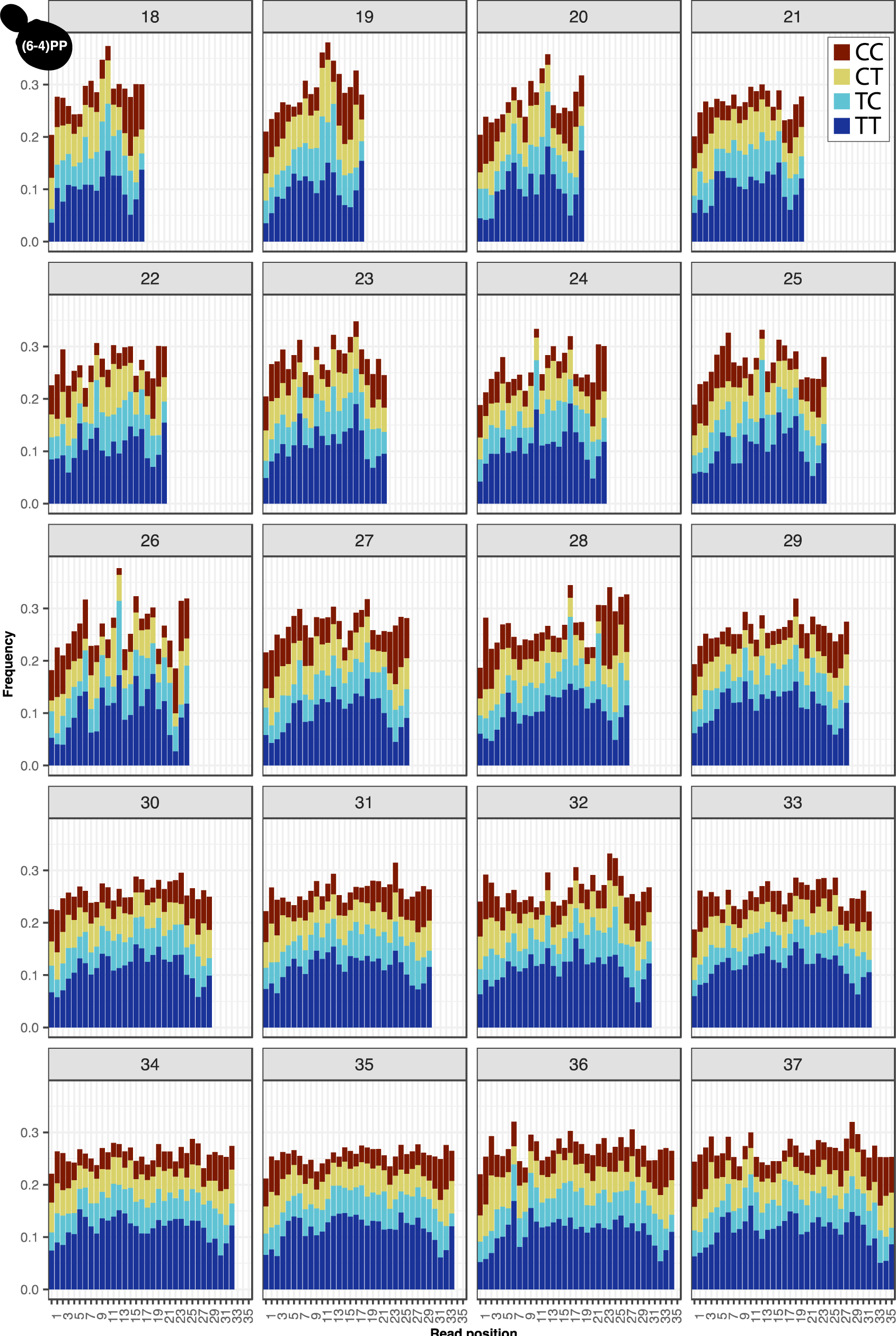
Di-pyrimidine frequencies in the mtDNA-mapping reads from the *S. cerevisiae* anti- (6-4)PP libraries for all read-length class.

**Figure S9.**
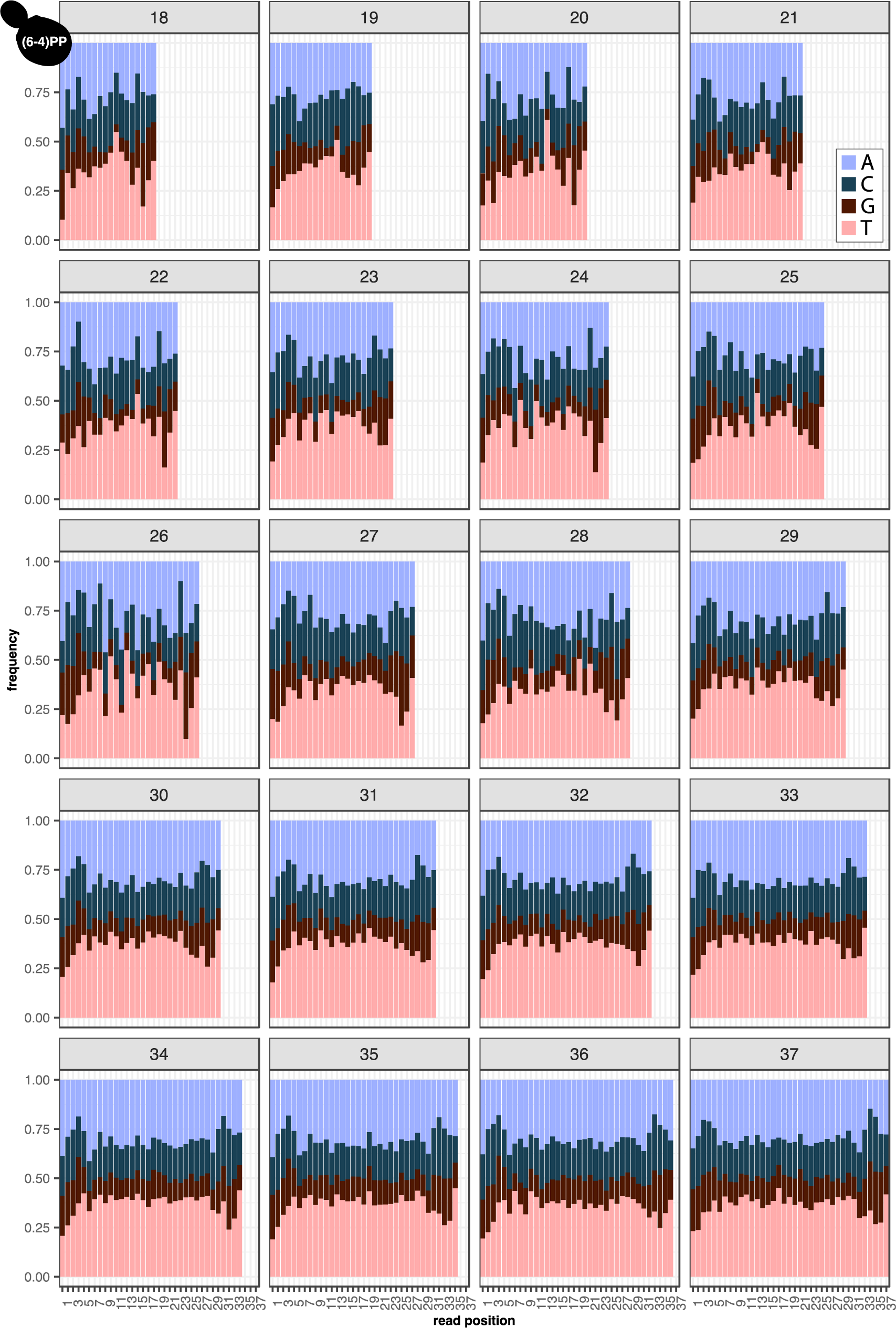
Nucleotide frequencies in the mtDNA-mapping reads from the *S. cerevisiae* anti-(6-4)PP libraries for all read-length classes.

**Figure S10.**
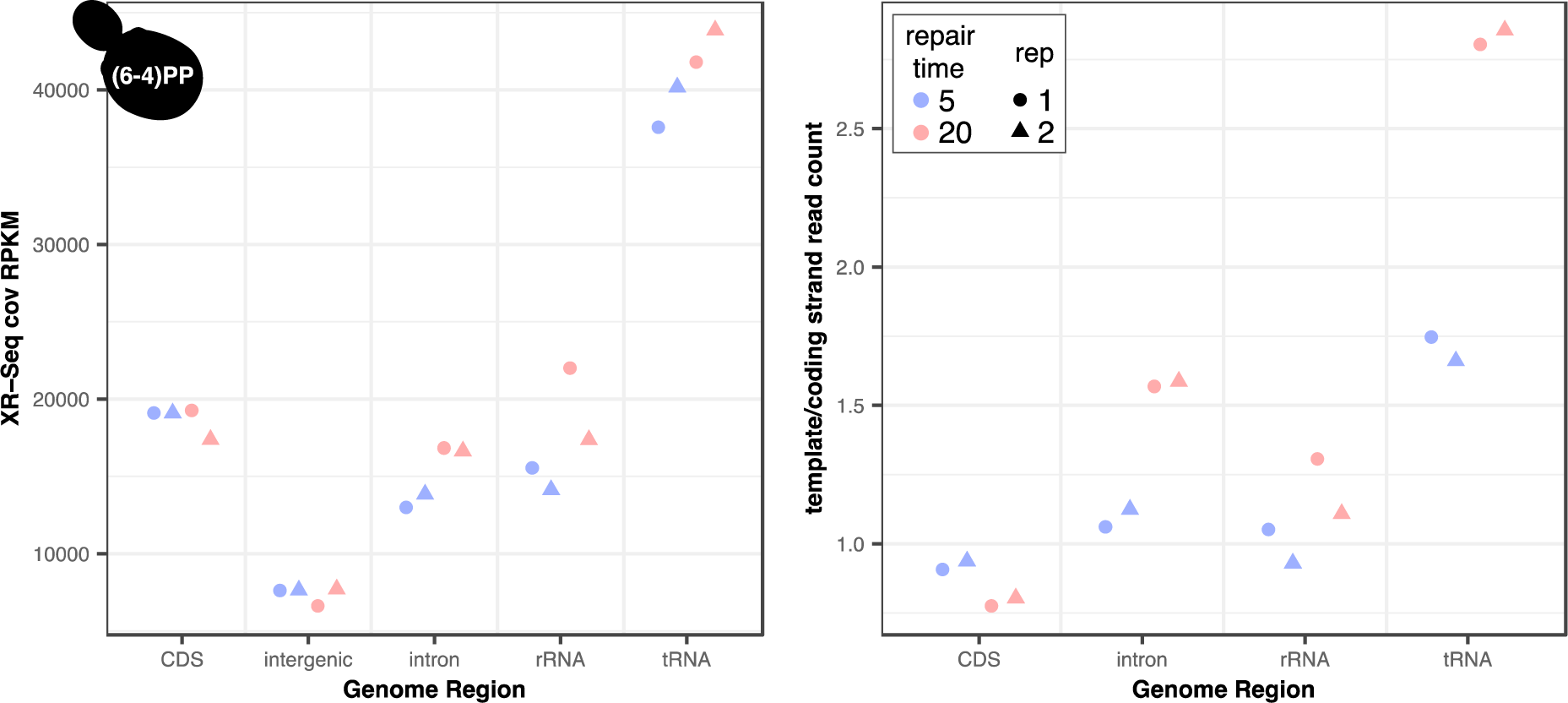
Depth of coverage (RPKM) and strand assymetry by genomic region in the mtDNA-mapping reads from the *S. cerevisiae* anti-(64)PP libraries.

**Figure S11.**
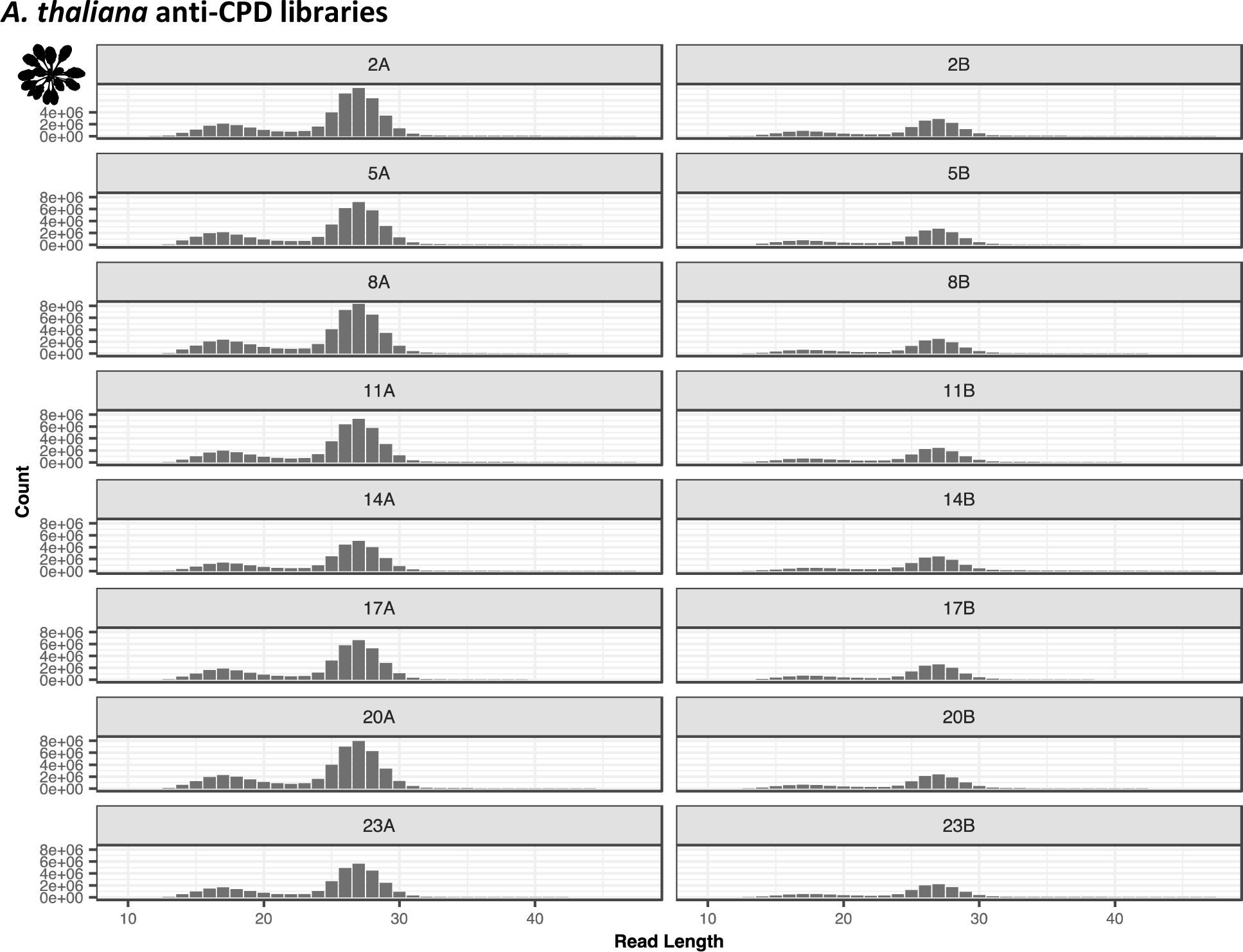
Length distributions of the nuclear-mapping reads from the *A. thaliana* anti-CPD libraries. The panel labels show the library ID, with the number indicating the time of day the plants were irradiated (2, 5, 8, 11, 14, 17, 20 or 23 hours) and the letter indicating the replicate (A or B).

**Figure S12.**
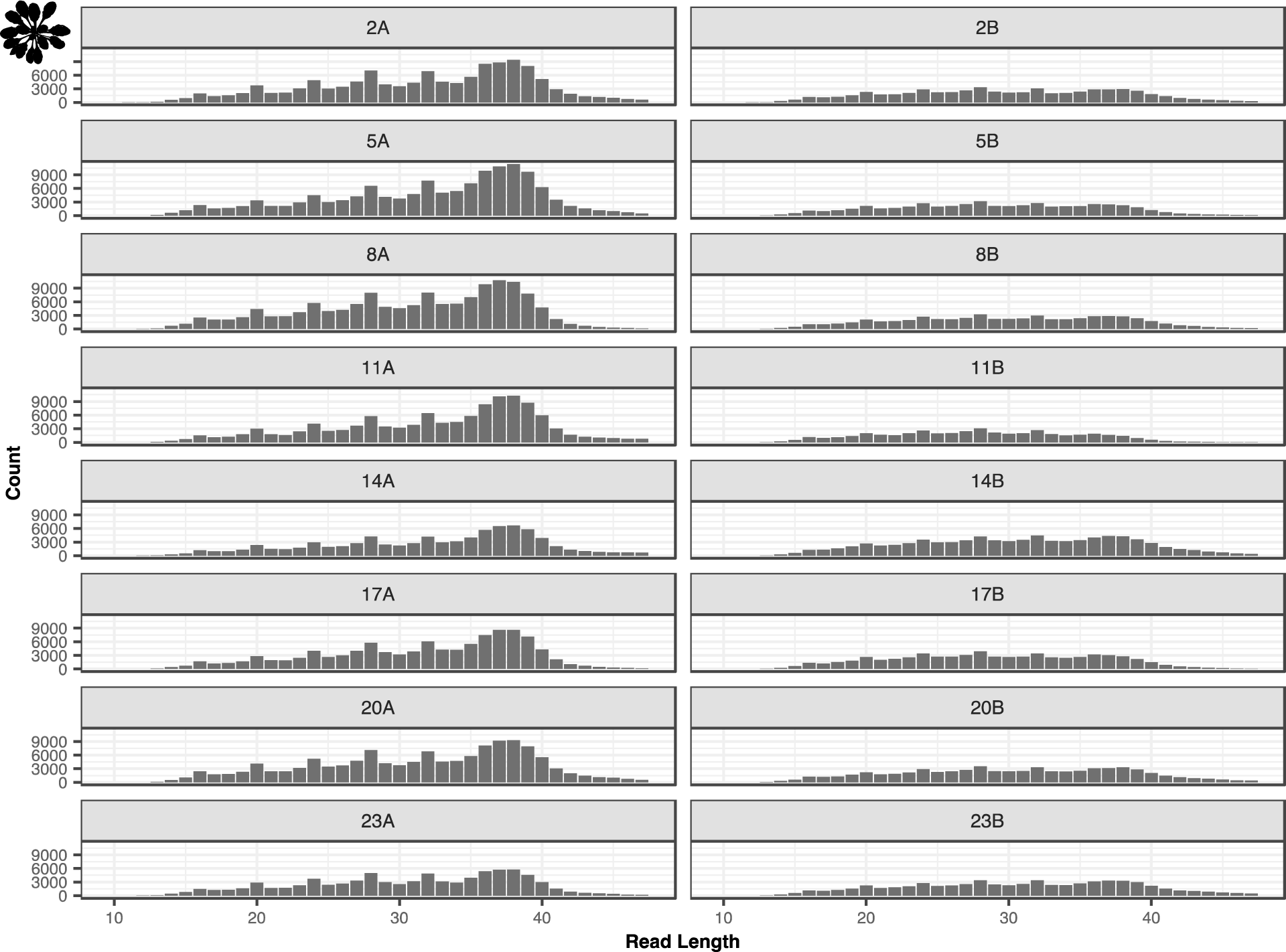
Length distributions of the mtDNA-mapping reads from the *A. thaliana* anti-CPD libraries. The panel labels show the library ID, with the number indicating the time of day the plants were irradiated (2, 5, 8, 11, 14, 17, 20 or 23 hours) and the letter indicating the replicate (A or B).

**Figure S13.**
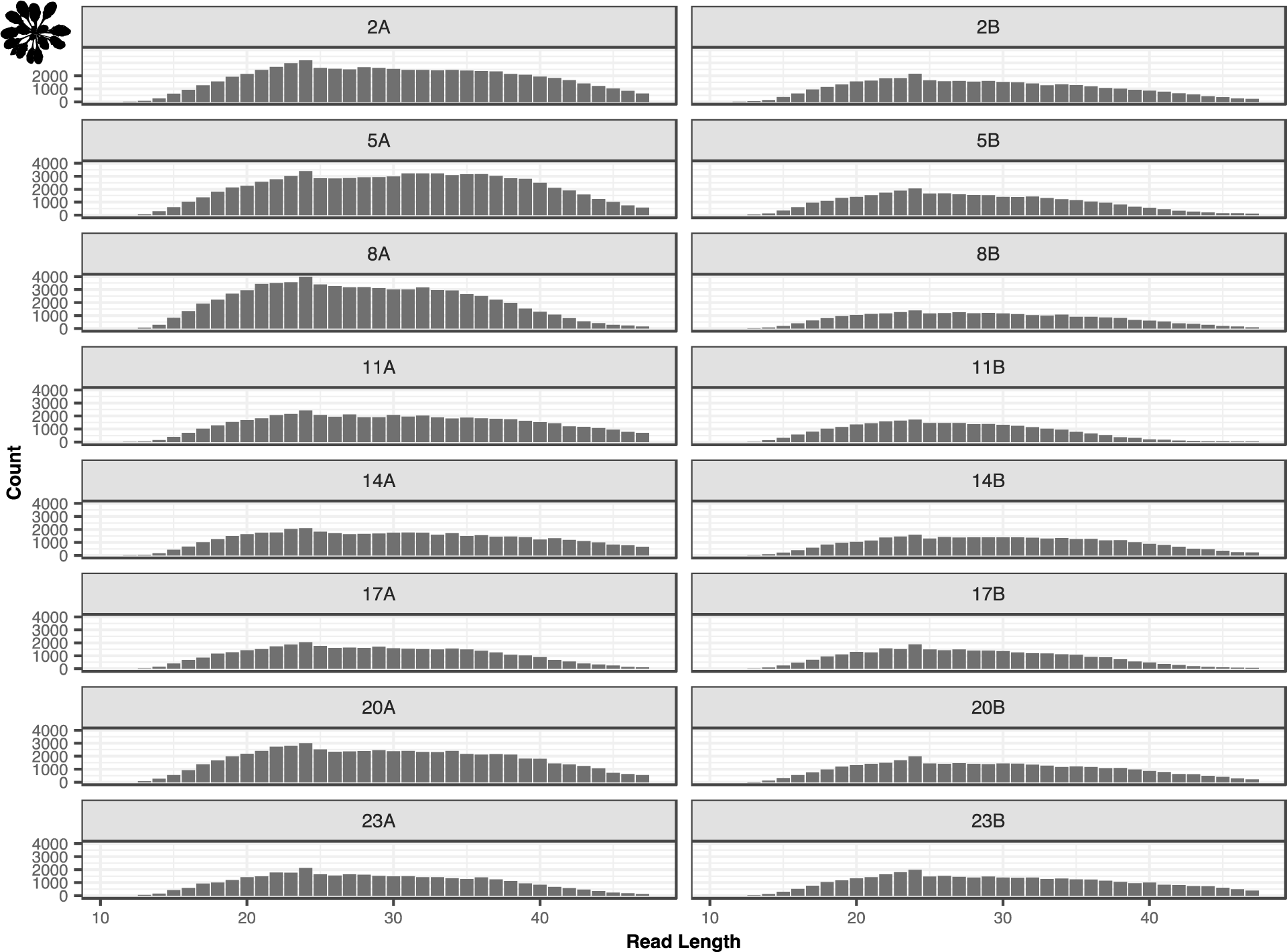
Length distributions of the ptDNA-mapping reads from the *A. thaliana* anti-CPD libraries. The panel labels show the library ID, with the number indicating the time of day the plants were irradiated (2, 5, 8, 11, 14, 17, 20 or 23 hours) and the letter indicating the replicate (A or B).

**Figure S14.**
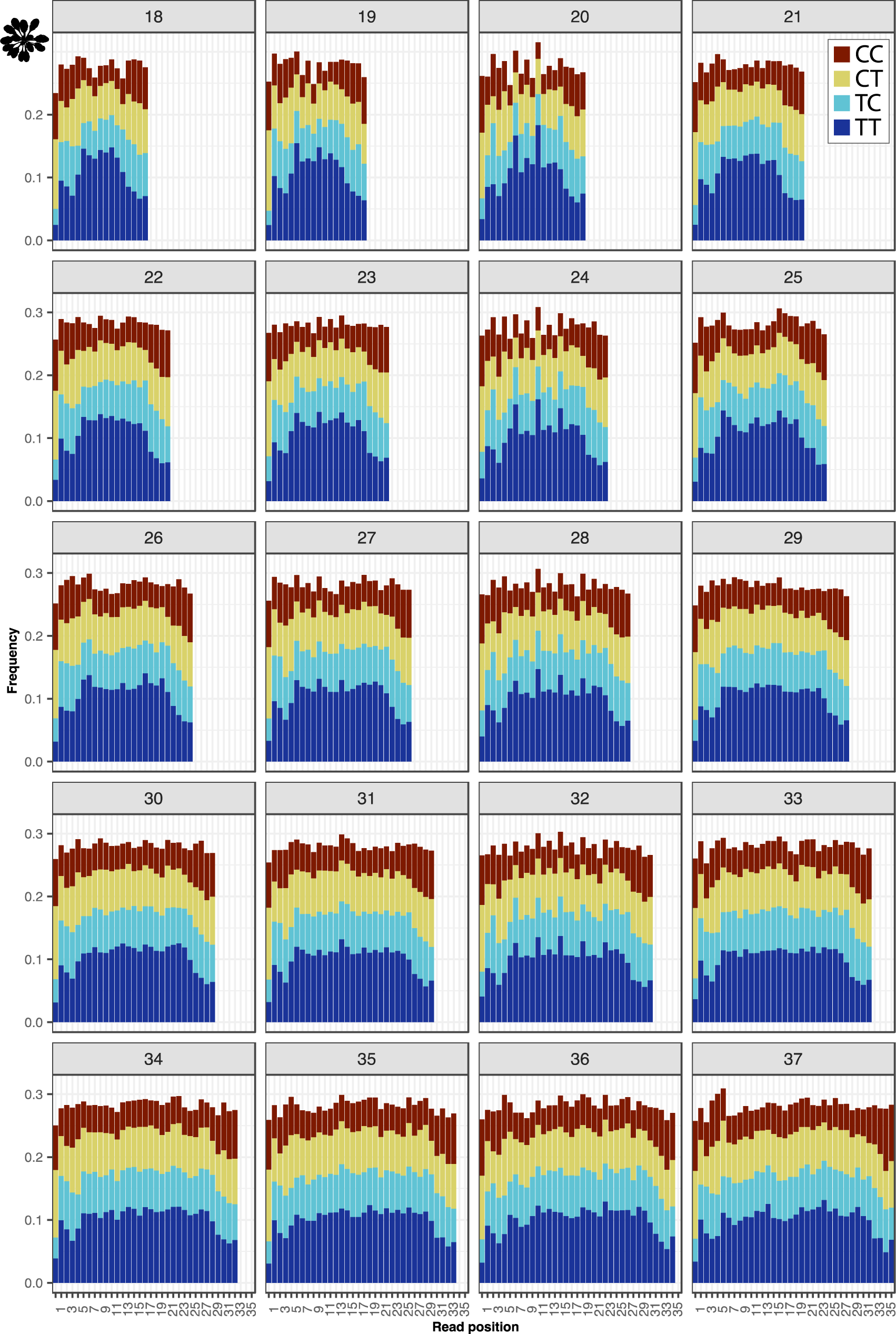
Di-pyrimidine frequencies in the mtDNA-mapping reads from the *A. thaliana* anti-CPD libraries.

**Figure S15.**
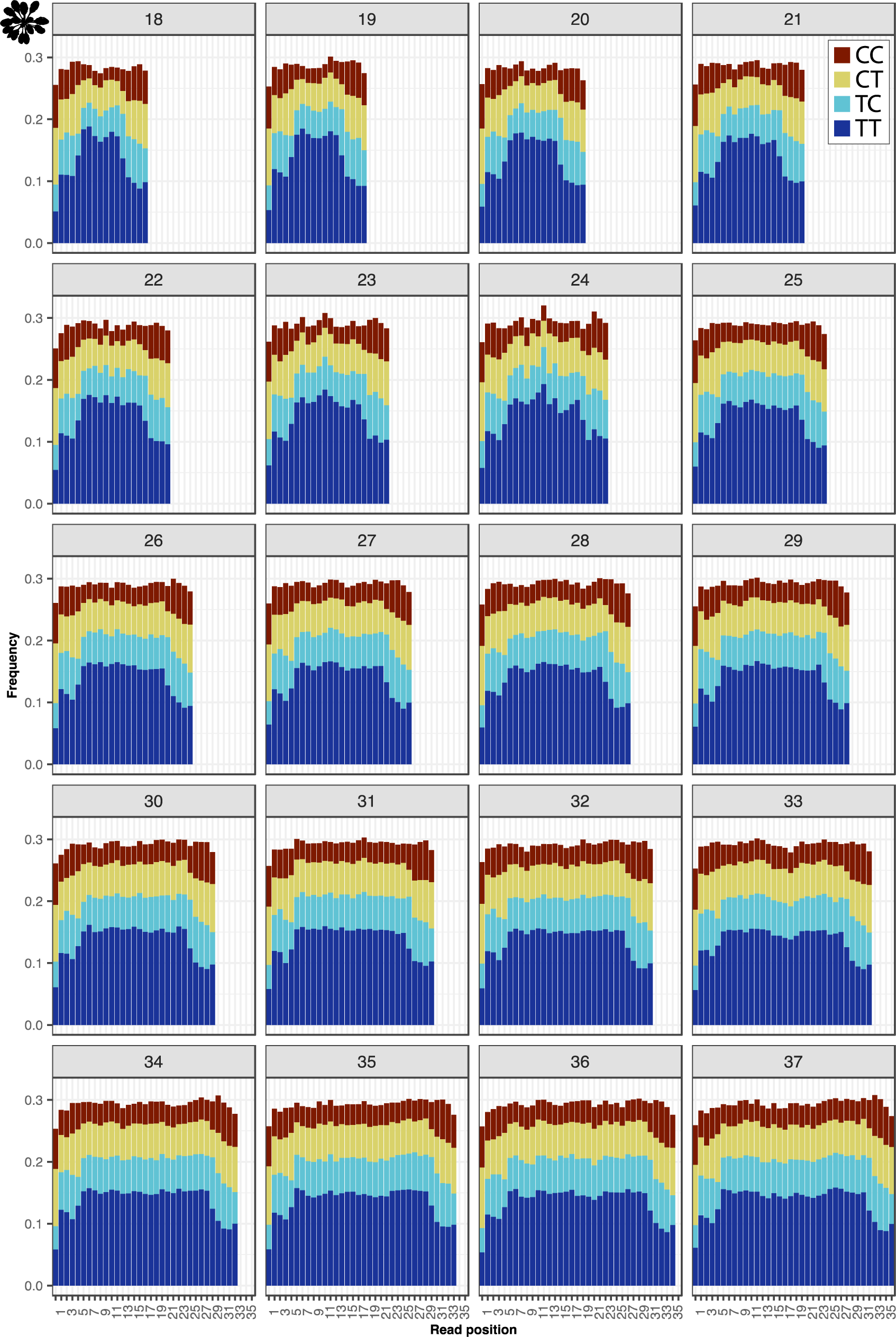
Di-pyrimidine frequencies in the ptDNA-mapping reads from the *A. thaliana* anti-CPD libraries.

**Figure S16.**
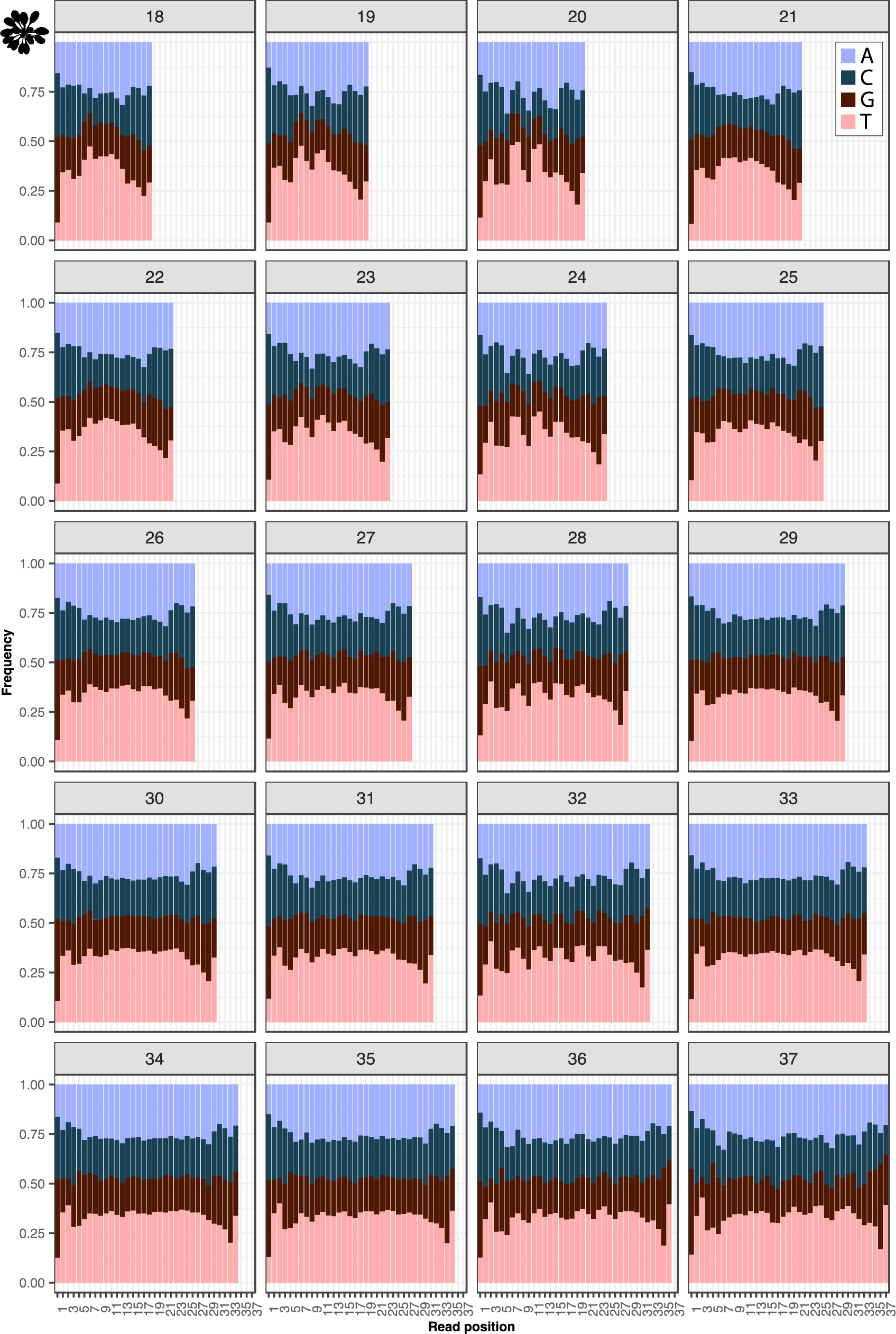
Nucleotide frequencies in the mtDNA-mapping reads from the *A. thaliana* anti-CPD libraries for all read-length classes.

**Figure S17.**
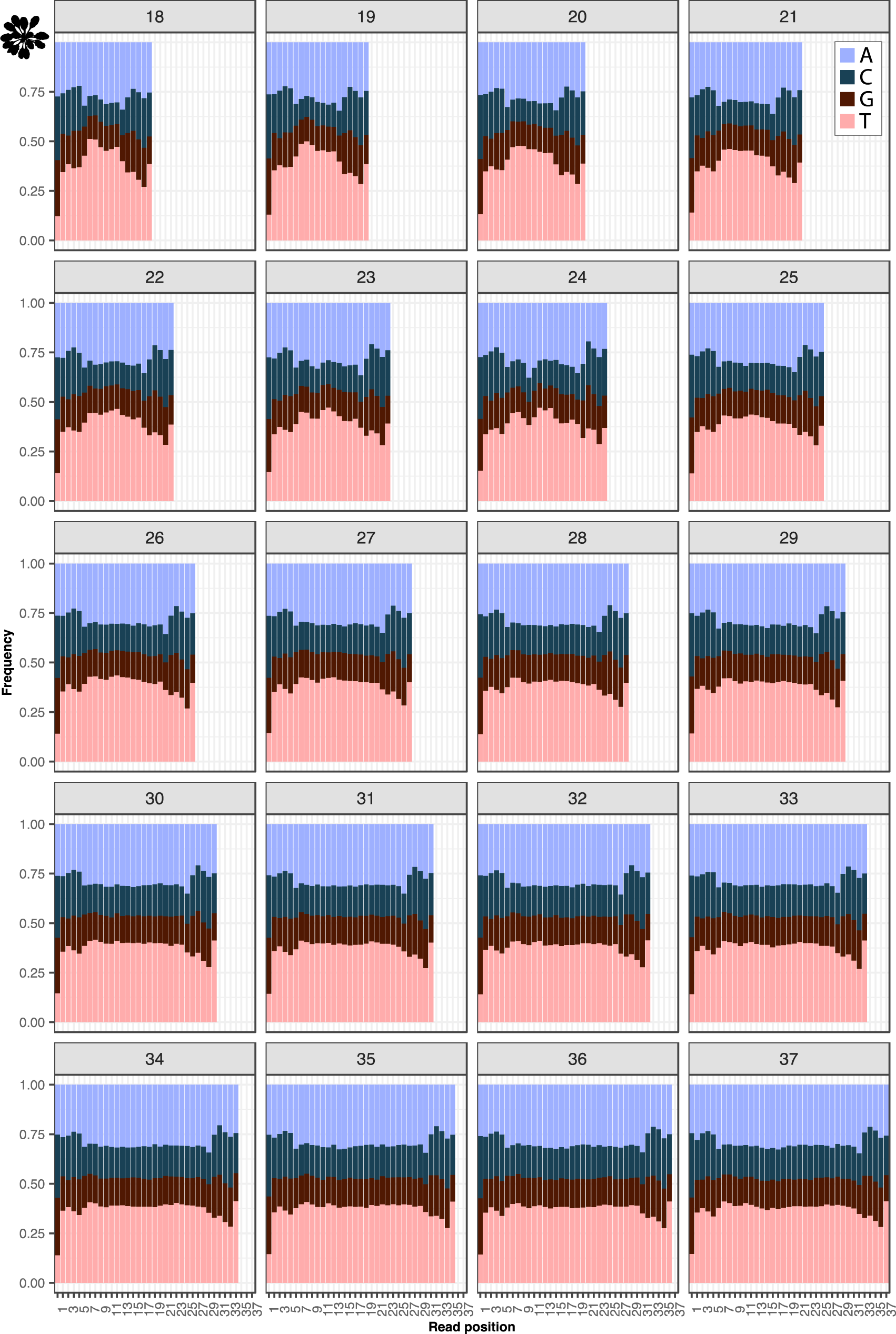
Nucleotide frequencies in the ptDNA-mapping reads from the *A. thaliana* anti-CPD libraries for all read-length classes.

**Figure S18.**
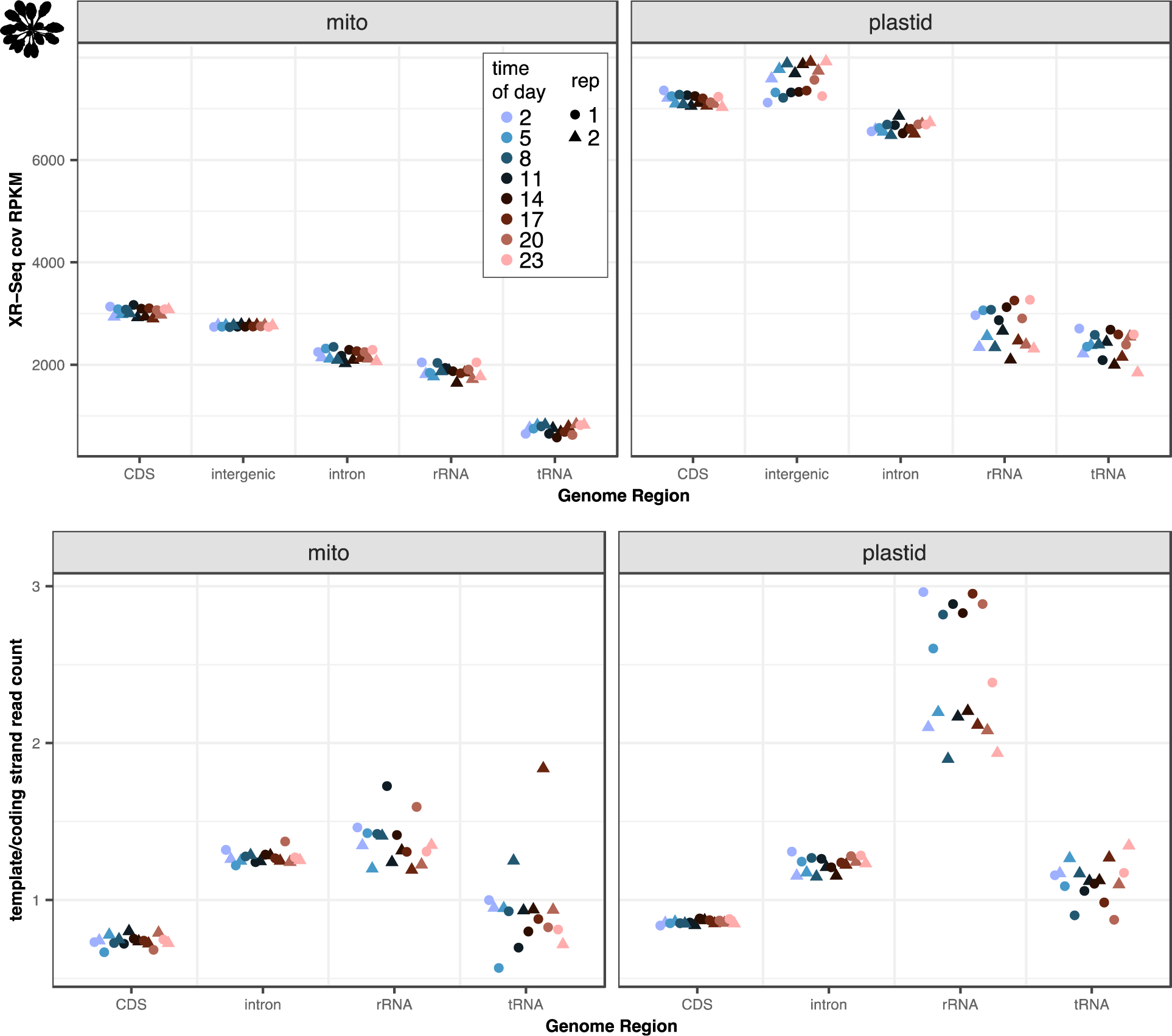
Depth of coverage (RPKM) and strand assymetry by genomic region in the mtDNA- and ptDNA-mapping reads from the *A. thaliana* anti-CPD libraries.

**Figure S19.**
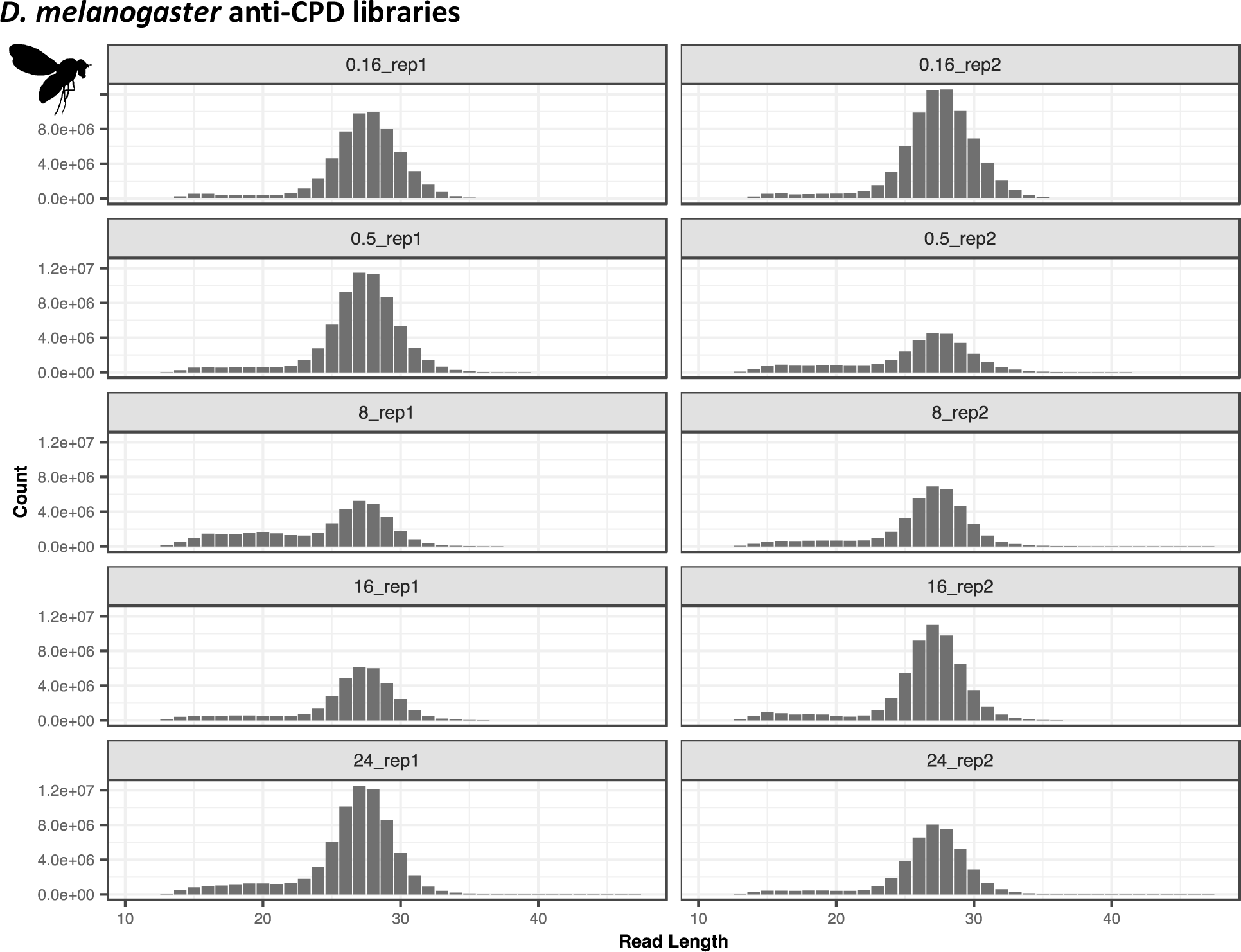
Length distributions of the nuclear-mapping reads from the *D. melanogaster* anti-CPD libraries. The panel labels show the library ID, with the number indicating the repair time (0.16, 0.5, 8, 16, or 24 hours) and the rep# indicating the replicate (1 or 2).

**Figure S20.**
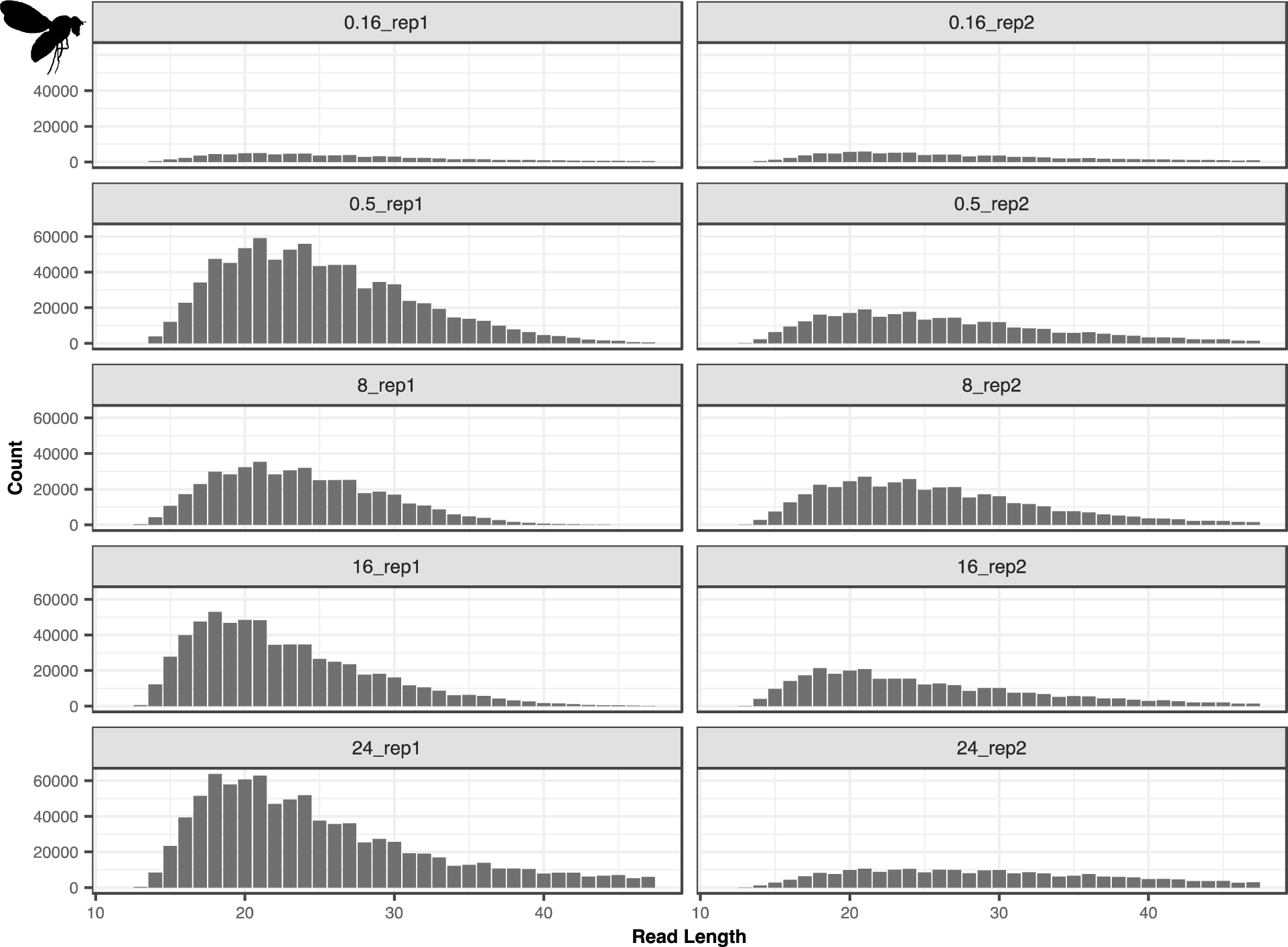
Length distributions of the mtDNA-mapping reads from the *D. melanogaster* anti-CPD libraries. The panel labels show the library ID, with the number indicating the repair time (0.16, 0.5, 8, 16, or 24 hours) and the rep# indicating the replicate (1 or 2).

**Figure S21.**
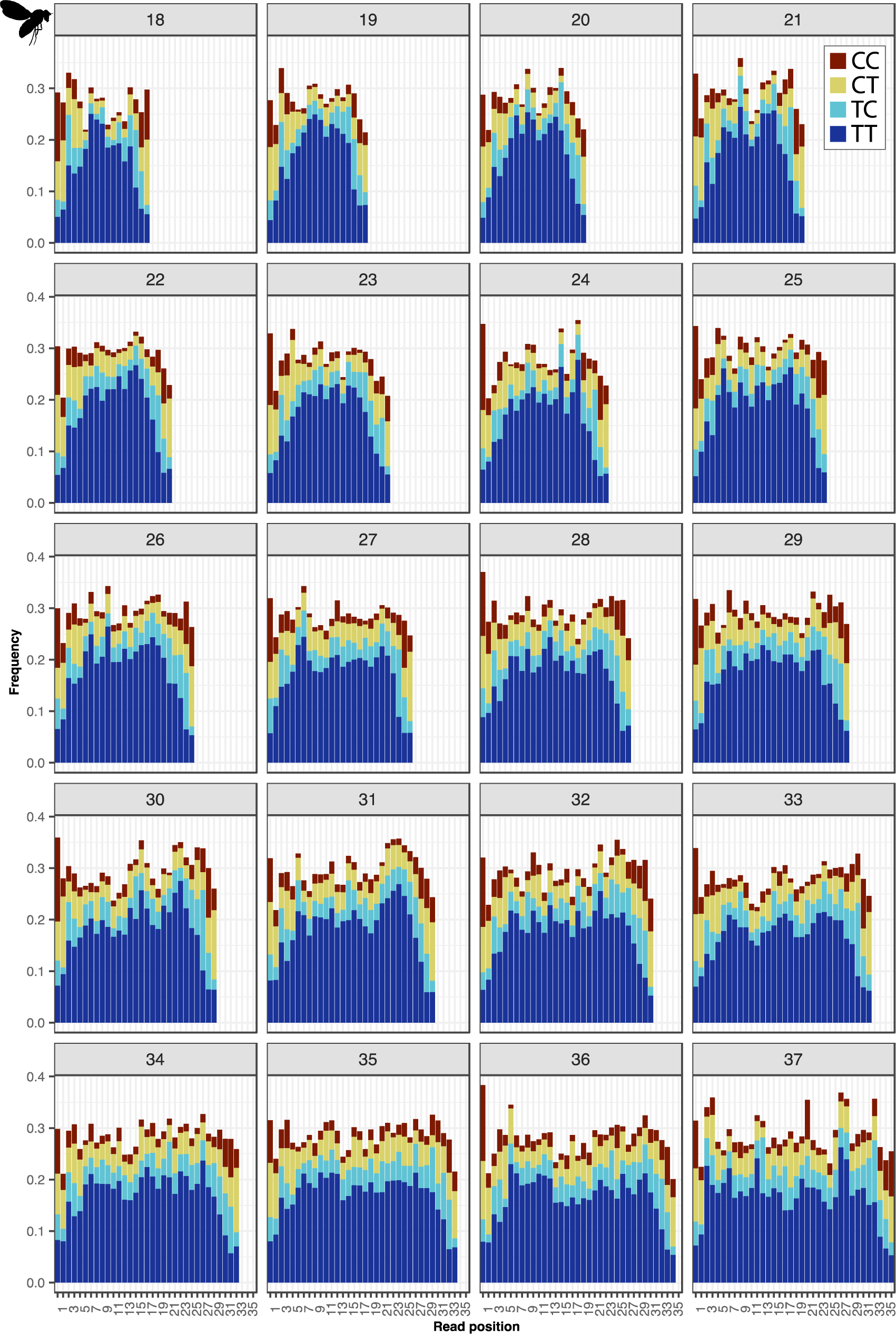
Di-pyrimidine frequencies in the mtDNA-mapping reads from the *D. melanogaster* anti-CPD libraries for all read-length classes.

**Figure S22.**
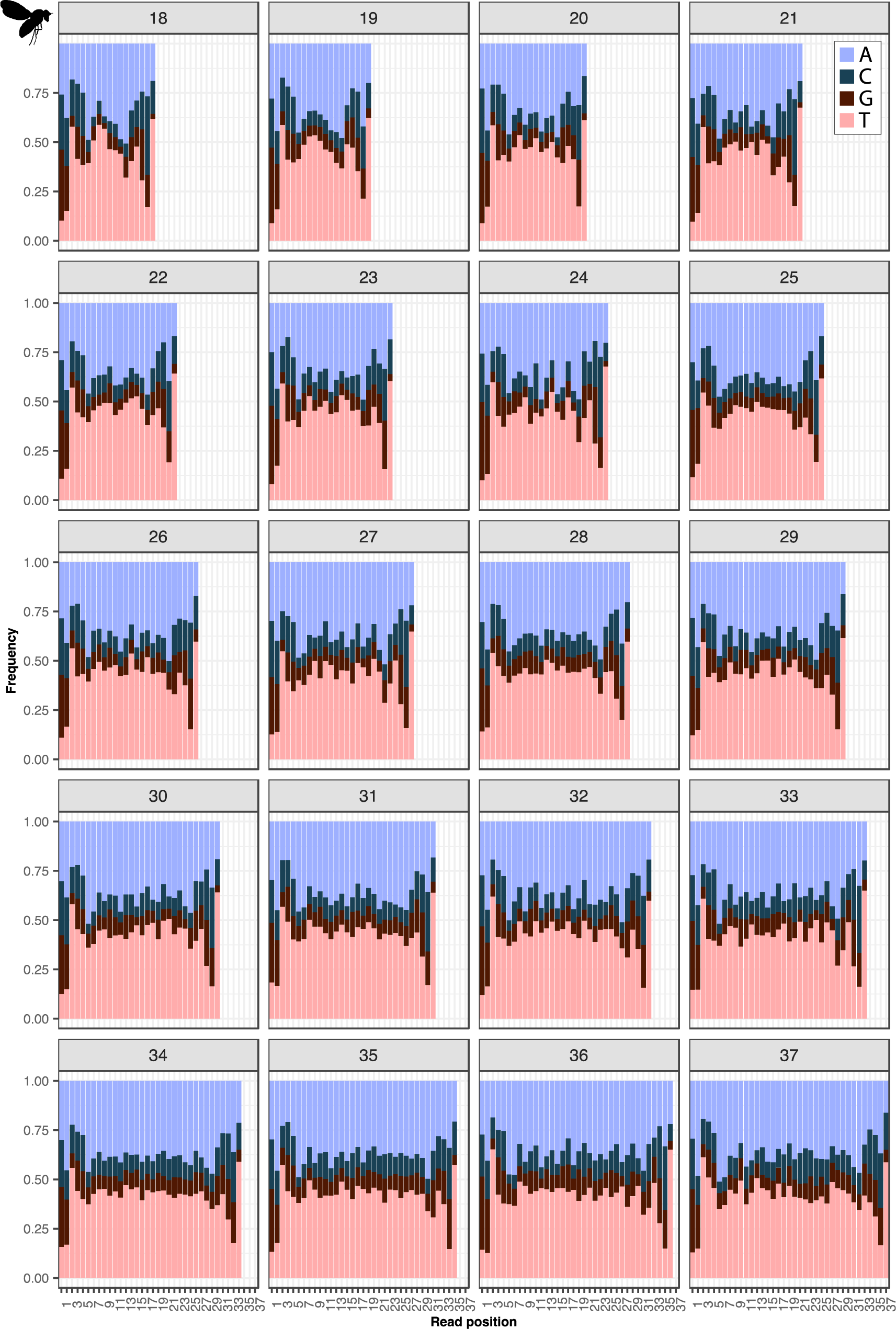
Nucleotide frequencies in the mtDNA mapping reads from the *D. melanogaster* anti-CPD libraries for all read-length classes.

**Figure S23.**
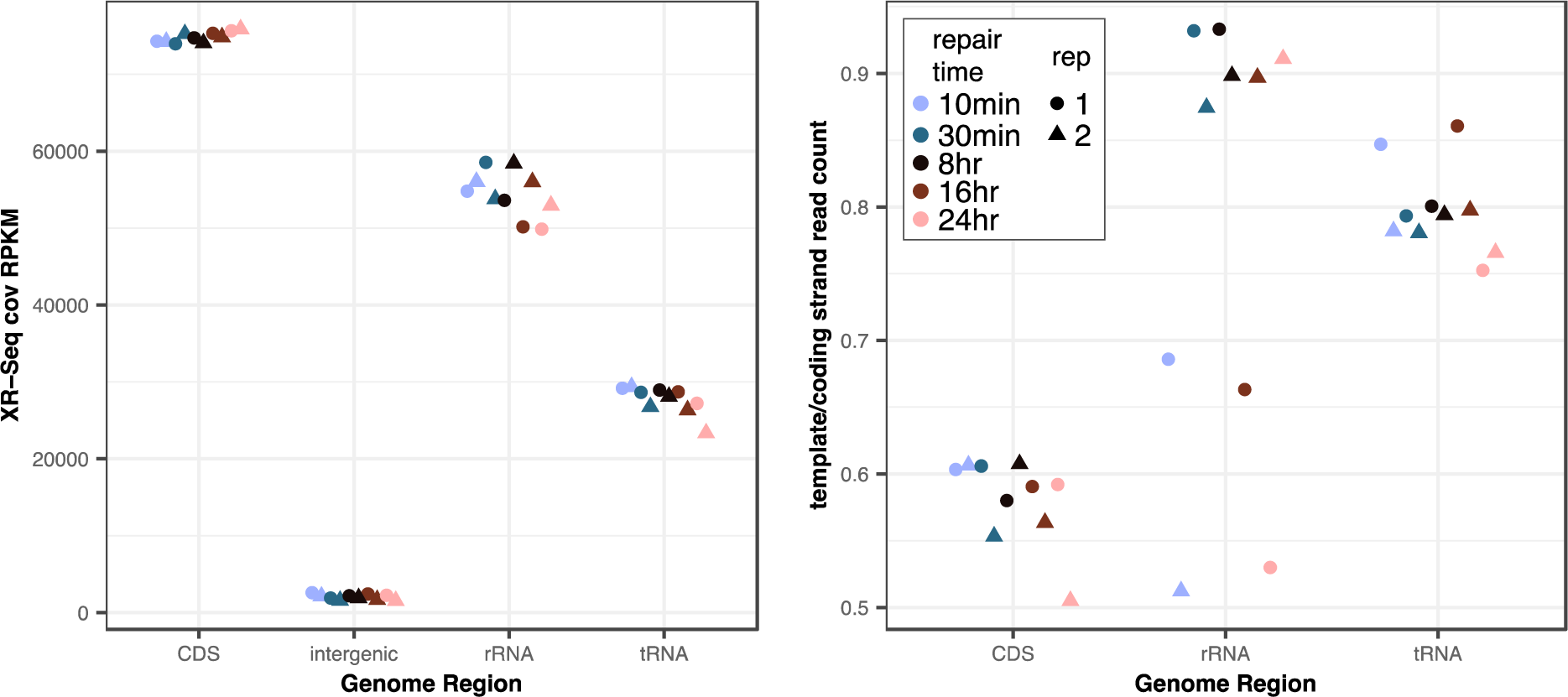
Depth of coverage (RPKM) and strand assymetry by genomic region in the mtDNA-mapping reads from the *D. melanogaster* anti-CPD libraries.

**Figure S24.**
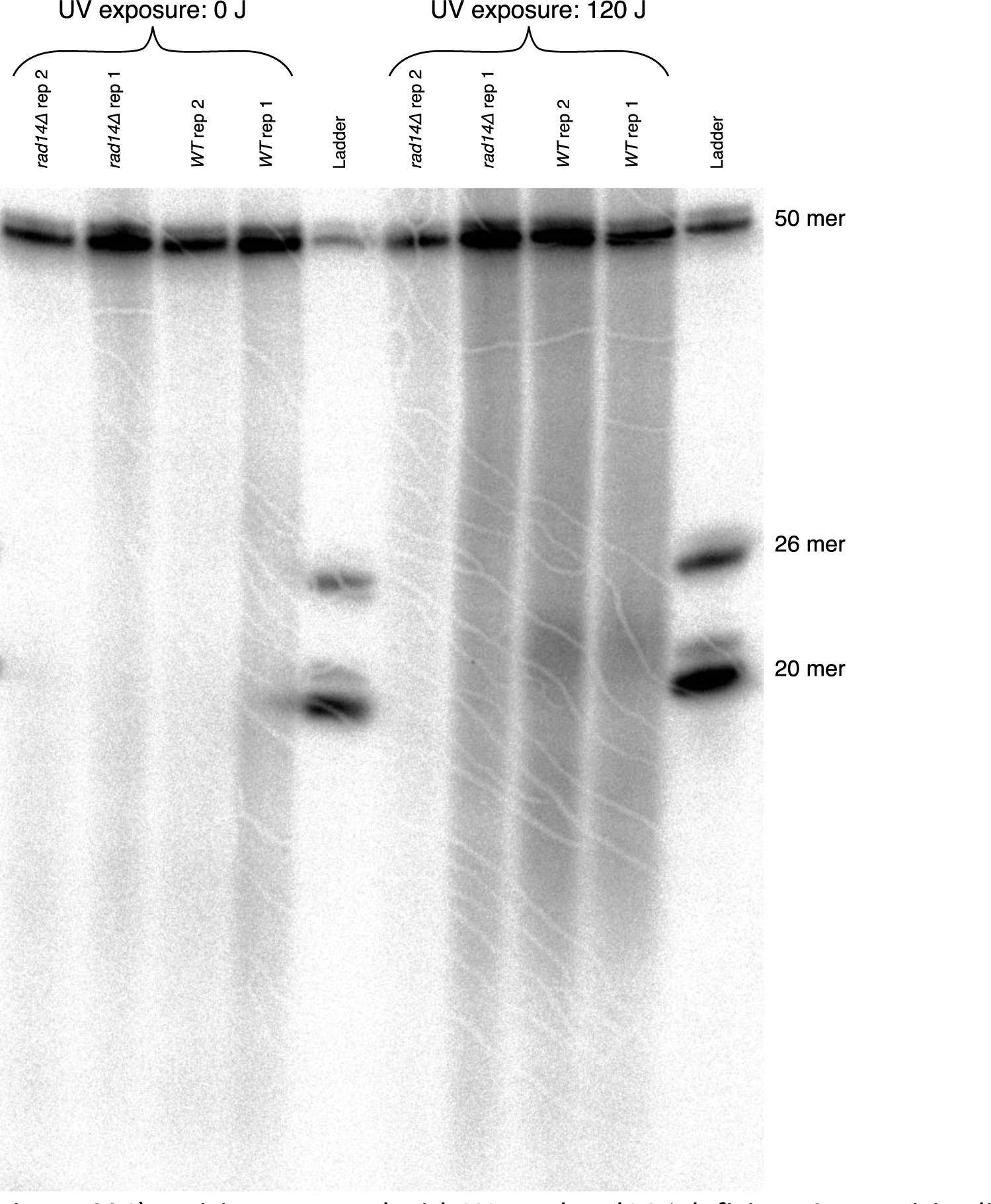
Excision assay gel with WT and *rad14Δ* deficient *S. cerevisiae* lines either not exposed to UV (left) or exposed to UV and given 20 minutes of dark repair (right). In conventional excision assays (50, 51), 25mer and 50mer PAGE purified oligos are radiolabeled alongside samples of interest to serve as a ladder in the high density polyacrylamide electrophoresis. Under our prediction that the nuclear-NER mutant would produce only mtDNA-derived fragments, we expected to detect the 26-nt, 24-nt and 22-nt peaks that punctuate the *S. cerevisiae* read length distribution (Figure 1), and thus used PAGE purified oligos of 50, 26, and 20 nt in length to make a custom ladder. Though our 11 % acrylamide gel cracked during drying, the samples and bands can still be reliably identified. In the UV-exposed WT samples, note the darker area between the 26mer and the 20mer markers, and the slightly less dark area below the 20mer marker, corresponding to the primary products and degraded products in the excsion assay shown in Figure 1 (and Figure 2 of (58)). We predicted to detect mtDNA-derived peaks at 26, 24 and 22 nt in the *rad14Δ* lanes, but such bands were not apparent, likely due the high background noise of this assay.

**Table S1:**
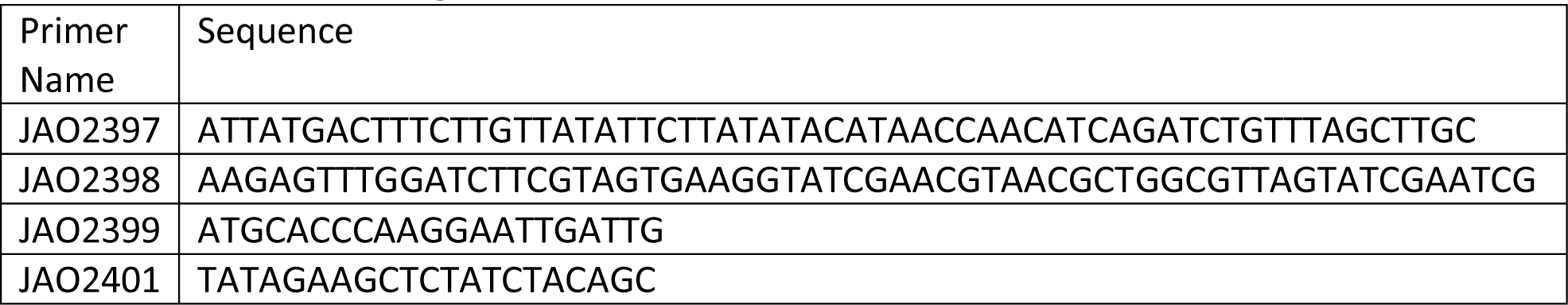
Primers used for generation and confirmation of *rad14Δ*.

## REFERENCES

1. Wolfe, K.H., Li, W.H. and Sharp, P.M. (1987) Rates of nucleotide substitution vary greatly among plant mitochondrial, chloroplast, and nuclear DNAs. Proc. Natl. Acad. Sci., 84, 9054–9058.

2. Drouin, G., Daoud, H. and Xia, J. (2008) Relative rates of synonymous substitutions in the mitochondrial, chloroplast and nuclear genomes of seed plants. Mol. Phylogenet. Evol., 49, 827–831.

3. Smith, D.R., Hua, J., Lee, R.W. and Keeling, P.J. (2012) Relative rates of evolution among the three genetic compartments of the red alga Porphyra differ from those of green plants and do not correlate with genome architecture. Mol. Phylogenet. Evol., 65, 339–344.

4. Havird, J.C. and Sloan, D.B. (2016) The roles of mutation, selection, and expression in determining relative rates of evolution in mitochondrial versus nuclear genomes. Mol. Biol. Evol., 33, 3042–3053.

5. Rong, Z., Tu, P., Xu, P., Sun, Y., Yu, F., Tu, N., Guo, L. and Yang, Y. (2021) The Mitochondrial Response to DNA Damage. Front. Cell Dev. Biol., 9, 1–10.

6. St. John, J.C. (2016) Mitochondrial DNA copy number and replication in reprogramming and differentiation. Semin. Cell Dev. Biol., 52, 93–101.

7. Clayton, D.A., Doda, J.N. and Friedberg, E.C. (1974) The absence of a pyrimidine dimer repair mechanism in mammalian mitochondria. Proc. Natl. Acad. Sci., 71, 2777–2781.

8. Druzhyna, N.M., Wilson, G.L. and LeDoux, S.P. (2008) Mitochondrial DNA repair in aging and disease. Mech. Ageing Dev., 129, 383–390.

9. Harman, D. (1972) The Biologic Clock: The Mitochondria? J. Am. Geriatr. Soc., 20, 145–147.

10. Murphy, M.P. (2009) How mitochondria produce reactive oxygen species. Biochem. J., 417, 1–13.

11. Saki, M. and Prakash, A. (2017) DNA damage related crosstalk between the nucleus and mitochondria. Free Radic. Biol. Med., 107, 216–227.

12. Muthye, V. and Lavrov, D. V. (2021) Multiple Losses of MSH1, Gain of mtMutS, and Other Changes in the MutS Family of DNA Repair Proteins in Animals. Genome Biol. Evol., 13, 1– 8.

13. Mower, J.P., Sloan, D.B. and Alverson, A.J. (2012) Plant mitochondrial genome diversity.

14. Zardoya, R. (2020) Recent advances in understanding mitochondrial genome diversity. F1000Research, 9.

15. Kazak, L., Reyes, A. and Holt, I.J. (2012) Minimizing the damage: Repair pathways keep mitochondrial DNA intact. Nat. Rev. Mol. Cell Biol., 13, 659–671.

16. Szczesny, B., Tann, A.W., Longley, M.J., Copeland, W.C. and Mitra, S. (2008) Long patch base excision repair in mammalian mitochondrial genomes. J. Biol. Chem., 283, 26349–26356.

17. Modrich, P. (2006) Mechanisms in eukaryotic mismatch repair. J. Biol. Chem., 281, 30305–30309.

18. de Souza-Pinto, N.C., Mason, P.A., Hashiguchi, K., Weissman, L., Tian, J., Guay, D., Lebel, M., Stevnsner, T. V., Rasmussen, L.J. and Bohr, V.A. (2009) Novel DNA mismatch-repair activity involving YB-1 in human mitochondria. DNA Repair (Amst)., 8, 704–719.

19. Wu, Z., Waneka, G., Broz, A.K., King, C.R. and Sloan, D.B. (2020) MSH1 is required for maintenance of the low mutation rates in plant mitochondrial and plastid genomes. Proc. Natl. Acad. Sci., 117, 16448–16455.

20. Wood, R.D. (1999) DNA damage recognition during nucleotide excision repair in mammalian cells. Biochimie, 81, 39–44.

21. Sancar, A. (1996) DNA excision repair. Annu. Rev. Biochem., 65, 43–81.

22. Prakash, L. (1975) Repair of pyrimidine dimers in nuclear and mitochondrial DNA of yeast irradiated with low doses of ultraviolet light. J. Mol. Biol., 98, 781–795.

23. Sakamoto, W. and Takami, T. (2018) Chloroplast DNA dynamics: Copy number, quality control and degradation. Plant Cell Physiol., 59, 1120–1127.

24. Zhao, L. (2019) Mitochondrial DNA degradation: A quality control measure for mitochondrial genome maintenance and stress response 1st ed. Elsevier Inc.

25. Bess, A.S., Ryde, I.T., Hinton, D.E. and Meyer, J.N. (2013) UVC-Induced Mitochondrial Degradation via Autophagy Correlates with mtDNA Damage Removal in Primary Human Fibroblasts. J. Biochem. Mol. Toxicol., 27, 28–41.

26. Bess, A.S., Crocker, T.L., Ryde, I.T. and Meyer, J.N. (2012) Mitochondrial dynamics and autophagy aid in removal of persistent mitochondrial DNA damage in Caenorhabditis elegans. Nucleic Acids Res., 40, 7916–7931.

27. Wang, H.T., Lin, J.H., Yang, C.H., Haung, C.H., Weng, C.W., Lin, A.M.Y., Lo, Y.L., Chen, W.S. and Tang, M.S. (2017) Acrolein induces mtDNA damages, mitochondrial fission and mitophagy in human lung cells. Oncotarget, 8, 70406–70421.

28. Dan, X., Babbar, M., Moore, A., Wechter, N., Tian, J., Mohanty, J.G., Croteau, D.L. and Bohr, V.A. (2020) DNA damage invokes mitophagy through a pathway involving Spata18. Nucleic Acids Res., 48, 6611–6623.

29. Shokolenko, I.N., Wilson, G.L. and Alexeyev, M.F. (2016) The ‘fast’ and the ‘slow’ modes of mitochondrial DNA degradation. Mitochondrial DNA, 27, 490–498.

30. Moretton, A., Morel, F., Macao, B., Lachaume, P., Ishak, L., Lefebvre, M., Garreau-Balandier, I., Vernet, P., Falkenberg, M. and Farge, G. (2017) Selective mitochondrial DNA degradation following double-strand breaks. PLoS One, 12, 1–17.

31. Urbina-Varela, R., Castillo, N., Videla, L.A. and Del Campo, A. (2020) Impact of mitophagy and mitochondrial unfolded protein response as new adaptive mechanisms underlying old pathologies: Sarcopenia and non-alcoholic fatty liver disease. Int. J. Mol. Sci., 21, 1–27.

32. Springer, M.Z. and Macleod, K.F. (2016) In Brief: Mitophagy: mechanisms and role in human disease. J. Pathol., 240, 253–255.

33. Doblado, L., Lueck, C., Rey, C., Samhan-arias, A.K., Prieto, I., Stacchiotti, A. and Monsalve, M. (2021) Mitophagy in human diseases. Int. J. Mol. Sci., 22, 1–51.

34. Emmerich, H.J., Saft, M., Schneider, L., Kock, D., Batschauer, A. and Essen, L.O. (2020) A topologically distinct class of photolyases specific for UV lesions within single-stranded DNA. Nucleic Acids Res., 48, 12845–12857.

35. Lucas-Lledó, J.I. and Lynch, M. (2009) Evolution of mutation rates: Phylogenomic analysis of the photolyase/cryptochrome family. Mol. Biol. Evol., 26, 1143–1153.

36. Goosen, N. and Moolenaar, G.F. (2008) Repair of UV damage in bacteria. DNA Repair (Amst)., 7, 353–379.

37. Mei, Q. and Dvornyk, V. (2015) Evolutionary history of the photolyase/cryptochrome superfamily in eukaryotes. PLoS One, 10, 1–20.

38. Takahashi, M., Teranishi, M., Ishida, H., Kawasaki, J., Takeuchi, A., Yamaya, T., Watanabe, M., Makino, A. and Hidema, J. (2011) Cyclobutane pyrimidine dimer (CPD) photolyase repairs ultraviolet-B-induced CPDs in rice chloroplast and mitochondrial DNA. Plant J., 66, 433– 442.

39. Yasui, A., Yajima, H., Kobayashi, T., Eker, A.P.M. and Oikawa, A. (1992) Mitochondrial DNA repair by photolyase. Mutat. Res. Repair, 273, 231–236.

40. Waters, R. and Moustacchi, E. (1974) The fate of ultraviolet-induced pyrimidine dimers in the mitochondrial DNA of Saccharomyces cerevisiae following various post-irradiation cell treatments. BBA Sect. Nucleic Acids Protein Synth., 366, 241–250.

41. Hunter, S.E., Jung, D., Di Giulio, R.T. and Meyer, J.N. (2010) The QPCR assay for analysis of mitochondrial DNA damage, repair, and relative copy number. Methods, 51, 444–451.

42. Ledoux, S.P., Wilson, G.L., Beecham, E.J., Stevnsner, T., Wassermann, K. and Bohr, V.A. (1992) Repair of mitochondrial DNA after various types of DNA damage in chinese hamster ovary cells. Carcinogenesis, 13, 1967–1973.

43. Kalinowski, D.P., Illenye, S. and van Houten, B. (1992) Analysis of DNA damage and repair in murine leukemia L1210 cells using a quantitative polymerase chain reaction assay. Nucleic Acids Res., 20, 3485–3494.

44. Sancar, A. (2004) Photolyase and cryptochrom blue-light photoreceptors. Adv. Protein Chem., 69, 73–100.

45. Mao, P., Smerdon, M.J., Roberts, S.A. and Wyrick, J.J. (2016) Chromosomal landscape of UV damage formation and repair at single-nucleotide resolution. Proc. Natl. Acad. Sci., 113, 9057–9062.

46. Hu, J., Adebali, O., Adar, S. and Sancar, A. (2017) Dynamic maps of UV damage formation and repair for the human genome. Proc. Natl. Acad. Sci., 114, 6758–6763.

47. Alhegaili, A.S., Ji, Y., Sylvius, N., Blades, M.J., Karbaschi, M., Tempest, H.G., Jones, G.D.D. and Cooke, M.S. (2019) Genome-wide adductomics analysis reveals heterogeneity in the induction and loss of cyclobutane thymine dimers across both the nuclear and mitochondrial genomes. Int. J. Mol. Sci., 20.

48. Hu, J., Adar, S., Selby, C.P., Lieb, J.D. and Sancar, A. (2015) Genome-wide analysis of human global and transcription-coupled excision repair of UV damage at single-nucleotide resolution. Genes Dev., 29, 948–960.

49. Amente, S., Di Palo, G., Scala, G., Castrignanò, T., Gorini, F., Cocozza, S., Moresano, A., Pucci, P., Ma, B., Stepanov, I., et al. (2019) Genome-wide mapping of 8-oxo-7,8-dihydro-2′-deoxyguanosine reveals accumulation of oxidatively-generated damage at DNA replication origins within transcribed long genes of mammalian cells. Nucleic Acids Res., 47, 221–236.

50. Hu, J., Choi, J.H., Gaddameedhi, S., Kemp, M.G., Reardon, J.T. and Sancar, A. (2013) Nucleotide excision repair in human cells: Fate of the excised oligonucleotide carrying dna damage in vivo. J. Biol. Chem., 288, 20918–20926.

51. Hu, J., Li, W., Adebali, O., Yang, Y., Oztas, O., Selby, C.P. and Sancar, A. (2019) Genome-wide mapping of nucleotide excision repair with XR-seq. Nat. Protoc., 14, 248–282f.

52. Mori, T., Nakane, M., Hattori, T., Matsunaga, T., Ihara, M. and Nikaido, O. (1991) Simultaneous Establishment of Monoclonal Antibodies Specific for Either Cyclobutane Pyrimidine Dimer or (6-4)Photoproduct From the Same Mouse Immunized With Ultraviolet-Irradiated Dna. Photochem. Photobiol., 54, 225–232.

53. Oztas, O., Selby, C.P., Sancar, A. and Adebali, O. (2018) Genome-wide excision repair in Arabidopsis is coupled to transcription and reflects circadian gene expression patterns. Nat. Commun., 9, 1–8.

54. Selby, C.P. and Sancar, A. (2006) A cryptochrome/photolyase class of enzymes with single-stranded DNA-specific photolyase activity. Proc. Natl. Acad. Sci., 103, 17696–17700.

55. Yang, Y., Adebali, O., Wu, G., Selby, C.P., Chiou, Y.Y., Rashid, N., Hu, J., Hogenesch, J.B. and Sancar, A. (2018) Cisplatin-DNA adduct repair of transcribed genes is controlled by two circadian programs in mouse tissues. Proc. Natl. Acad. Sci., 115, E4777–E4785.

56. Akkose, U., Kaya, V.O., Lindsey-Boltz, L., Karagoz, Z., Brown, A.D., Larsen, P.A., Yoder, A.D., Sancar, A. and Adebali, O. (2020) Comparative analyses of two primate species diverged by more than 60 million years show different rates but similar distribution of genome-wide UV repair events. BMC Genomics, 22, 1–13.

57. Deger, N., Yang, Y., Lindsey-Boltz, L.A., Sancar, A. and Selby, C.P. (2019) Drosophila, which lacks canonical transcription-coupled repair proteins, performs transcription-coupled repair. J. Biol. Chem., 294, 18092–18098.

58. Li, W., Adebali, O., Yang, Y., Selby, C.P. and Sancar, A. (2018) Single-nucleotide resolution dynamic repair maps of UV damage in Saccharomyces cerevisiae genome. Proc. Natl. Acad. Sci., 115, E3408–E3415.

59. Zhao, L. and Sumberaz, P. (2020) Mitochondrial DNA Damage: Prevalence, Biological Consequence, and Emerging Pathways. Chem. Res. Toxicol., 33, 2491–2502.

60. Lindsey-Boltz, L.A., Yang, Y., Kose, C., Deger, N., Eynullazada, K., Kawara, H. and Sancar, A. (2023) Nucleotide excision repair in Human cell lines lacking both XPC and CSB proteins. Nucleic Acids Res., 51, 6238–6245.

61. Andrews, S. (2010) Babraham bioinformatics-FastQC a quality control tool for high throughput sequence data. *URL https//www. bioinformatics. babraham. ac. uk/projects/fastqc*.

62. Martin, M. (2011) Cutadapt removes adapter sequences from high-throuoghput sequencing reads. EMBnet.jouonal, 17, 10–12.

63. Langmead, B. and Salzberg, S.L. (2012) Fast gapped-read alignment with Bowtie 2. Nat. Methods, 9, 357–359.

64. Li, H., Handsaker, B., Wysoker, A., Fennell, T., Ruan, J., Homer, N., Marth, G., Abecasis, G. and Durbin, R. (2009) The Sequence Alignment/Map format and SAMtools. Bioinformatics, 25, 2078–2079.

65. Fields, P.D., Waneka, G., Naish, M., Schatz, M.C., Henderson, I.R. and Sloan, D.B. (2022) Complete Sequence of a 641-kb Insertion of Mitochondrial DNA in the Arabidopsis thaliana Nuclear Genome. Genome Biol. Evol., 14, 1–9.

66. Guzder, S.N., Sommers, C.H., Prakash, L. and Prakash, S. (2006) Complex Formation with Damage Recognition Protein Rad14 Is Essential for Saccharomyces cerevisiae Rad1-Rad10 Nuclease To Perform Its Function in Nucleotide Excision Repair In Vivo. Mol. Cell. Biol., 26, 1135–1141.

67. Goldstein, A.L. and McCusker, J.H. (1999) Three new dominant drug resistance cassettes for gene disruption in Saccharomyces cerevisiae. Yeast, 15, 1541–1553.

68. Winston, F., Dollard, C. and Ricupero-Hovasse, S.L. (1995) Construction of a set of convenient saccharomyces cerevisiae strains that are isogenic to S288C. Yeast, 11, 53–55.

69. Boulé, J.B., Rougeon, F. and Papanicolaou, C. (2001) Terminal Deoxynucleotidyl Transferase Indiscriminately Incorporates Ribonucleotides and Deoxyribonucleotides. J. Biol. Chem., 276, 31388–31393.

70. Altschuler, S.E., Lewis, K.A. and Wuttke, D.S. (2013) Practical strategies for the evaluation of high-affinity protein/nucleic acid interactions. J. Nucleic Acids Investig., 4, 3.

71. Shen, J., Zhang, Y., Havey, M.J. and Shou, W. (2019) Copy numbers of mitochondrial genes change during melon leaf development and are lower than the numbers of mitochondria. Hortic. Res., 6.

72. O’Hara, R., Tedone, E., Ludlow, A., Huang, E., Arosio, B., Mari, D. and Shay, J.W. (2019) Quantitative mitochondrial DNA copy number determination using droplet digital PCR with single-cell resolution. Genome Res., 29, 1878–1888.

73. Herbers, E., Kekäläinen, N.J., Hangas, A., Pohjoismäki, J.L. and Goffart, S. (2019) Tissue specific differences in mitochondrial DNA maintenance and expression. Mitochondrion, 44, 85–92.

74. Göke, A., Schrott, S., Mizrak, A., Belyy, V., Osman, C. and Waltera, P. (2020) Mrx6 regulates mitochondrial DNA copy number in Saccharomyces cerevisiae by engaging the evolutionarily conserved Lon protease Pim1. Mol. Biol. Cell, 31, 527–545.

75. Gonzalez-Hunt, C.P., Wadhwa, M. and Sanders, L.H. (2018) DNA damage by oxidative stress: Measurement strategies for two genomes. Curr. Opin. Toxicol., 7, 87–94.

76. Adebali, O., Sancar, A. and Selby, C.P. (2017) Mfd translocase is necessary and sufficient for Transcription-coupled repair in Escherichia coli. J. Biol. Chem., 292, 18386–18391.

77. Adebali, O., Chiou, Y.Y., Hu, J., Sancar, A. and Selby, C.P. (2017) Genome-wide transcription-coupled repair in Escherichia coli is mediated by the Mfd translocase. Proc. Natl. Acad. Sci., 114, E2116–E2125.

78. Kim, S.H., Kim, G.H., Kemp, M.G. and Choi, J.H. (2022) TREX1 degrades the 3′ end of the small DNA oligonucleotide products of nucleotide excision repair in human cells. Nucleic Acids Res., 50, 3974–3984.

79. Kemp, M.G. and Sancar, A. (2012) DNA excision repair: Where do all the dimers go? Cell Cycle, 11, 2997–3002.

80. Li, W., Hu, J., Adebali, O., Adar, S., Yang, Y., Chiou, Y.Y. and Sancar, A. (2017) Human genome-wide repair map of DNA damage caused by the cigarette smoke carcinogen benzo[a]pyrene. Proc. Natl. Acad. Sci., 114, 6752–6757.

81. Bogenhagen, D.F. (2012) Mitochondrial DNA nucleoid structure. Biochim. Biophys. Acta - Gene Regul. Mech., 1819, 914–920.

82. Di Giorgio, J.A.P., Lepage, É., Tremblay-Belzile, S., Truche, S., Loubert-Hudon, A. and Brisson, N. (2019) Transcription is a major driving force for plastid genome instability in Arabidopsis. PLoS One, 14, 1–30.

83. Belozerova, N.S., Pozhidaeva, E.S., Shugaev, A.G. and Kusnetsov, V. V. (2011) Run-on transcription as a method for the analysis of mitochondrial genome expression. Russ. J. Plant Physiol., 58, 164–168.

84. Waneka, G., Svendsen, J.M., Havird, J.C. and Sloan, D.B. (2021) Mitochondrial mutations in Caenorhabditis elegans show signatures of oxidative damage and an AT-bias . Genetics, 219.

85. Wu, Z., Waneka, G. and Sloan, D.B. (2020) The tempo and mode of angiosperm mitochondrial genome divergence inferred from intraspecific variation in arabidopsis thaliana. G3 Genes, Genomes, Genet., 10, 1077–1086.

86. Daniel, A., Michael, R., Chen Wei-Sheng, Danielsson Maxwell, Fennell Timothy, Russ Carsten, Jaffe David, Nusbaum Chad and Andreas, G. (2011) Analyzing and minimizing PCR amplification bias in Illumina sequencing libraries. Genome Biol., 12, 1–14.

87. Stein, A. and Sia, E.A. (2017) Mitochondrial DNA repair and damage tolerance. Front. Biosci. - Landmark, 22, 920–943.

88. Chevigny, N., Schatz-Daas, D., Lotfi, F. and Gualberto, J.M. (2020) DNA repair and the stability of the plant mitochondrial genome. Int. J. Mol. Sci., 21.

89. Alencar, R.R., Batalha, C.M.P.F., Freire, T.S. and de Souza-Pinto, N.C. (2019) Enzymology of mitochondrial DNA repair 1st ed. Elsevier Inc.

90. Matic, S., Jiang, M., Nicholls, T.J., Uhler, J.P., Dirksen-Schwanenland, C., Polosa, P.L., Simard, M.L., Li, X., Atanassov, I., Rackham, O., et al. (2018) Mice lacking the mitochondrial exonuclease MGME1 accumulate mtDNA deletions without developing progeria. Nat. Commun., 9.

91. Peeva, V., Blei, D., Trombly, G., Corsi, S., Szukszto, M.J., Rebelo-Guiomar, P., Gammage, P.A., Kudin, A.P., Becker, C., Altmüller, J., et al. (2018) Linear mitochondrial DNA is rapidly degraded by components of the replication machinery. Nat. Commun., 9, 1–11.

